# Inhibition of METTL3 by STC-15 induces RNA misprocessing that results in dsRNA formation and activates innate immunity

**DOI:** 10.64898/2025.12.05.692335

**Authors:** Harry Fischl, Tamsin J Samuels, Byron Andrews, Lina Vasiliauskaitė, Daniel Jupp, Claire J Saunders, Joanna Obacz, Yaara Ofir-Rosenfeld, Oliver Rausch

## Abstract

The RNA methyltransferase METTL3 is responsible for the generation of m6A, the most abundant modification mark on mRNA and long non-coding RNA. Accumulating evidence suggests numerous roles of METTL3 in cancer initiation and progression and highlights the potential for targeting this enzyme in oncology. STC-15 is a potent and selective METTL3 inhibitor and the first RNA modifying enzyme inhibitor to enter human clinical development. It is structurally related to the previously published tool inhibitors STM2457 and STM3675.

We previously identified the induction of a cancer cell-intrinsic interferon response following pharmacological inhibition of METTL3, leading to activation of T-cell-mediated anti-tumour response. Here, we profiled m6A levels at nucleotide resolution using GLORI and characterised RNA changes following METTL3 inhibition with STC-15 or STM3675. Following loss of m6A, we uncovered aberrant mRNA transcripts arising from intron retention (IR) and transcriptional run-on (RO) events downstream of m6A-enriched exons in human cancer cells and in tumour samples *in vivo*. We found that these IR and RO events produce double-stranded RNA and are bound by the cytoplasmic dsRNA sensor MDA5. Using preclinical *in vitro* and *in vivo* models, we characterised in detail the anti-tumour immune responses induced by STC-15. Our study reveals how METTL3 inhibition leads to dsRNA accumulation, which triggers a type I interferon response and induces anti-tumour immunity. Together, these findings provide a mechanistic rationale for STC-15 as a novel anti-cancer drug both as monotherapy and in combination with anti-PD1 checkpoint inhibitors.

## Introduction

N⁶-Methyladenosine, or m6A, is the most abundant modification on mRNA and long non-coding RNA (lncRNA). It is generated by the RNA methyltransferase METTL3, which co-transcriptionally methylates the N6-position of selected adenosines. METTL3 is the active subunit of a heterodimer composed of METTL3 and METTL14 polypeptides. The heterodimer, in turn, is part of the bigger m6A writer complex. m6A modification directly affects transcript fate, by modulating fundamental processes such as splicing, localisation, stability and mRNA translation (He & He, 2021, Zaccara et al., 2019). The DRACH consensus motif has been identified for METTL3-dependent m6A modifications on mRNA (where D = A, G, or U; R = G or A; H = A, C or U) (He & He, 2021, Zaccara et al., 2019, Tzelepis et al., 2019). m6A modifications are not distributed uniformly along mRNA transcripts. Although they can be found anywhere along the transcript, mapping m6A sites revealed a strong enrichment near the stop codon (or near the last exon-exon junction), and in long internal exons (He & He, 2021, Koh et al., 2019).

Accumulating evidence suggests that RNA modifications are involved in cancer initiation and progression, and cancer cell-intrinsic roles for METTL3 have been identified in numerous cancers. For example, in AML, METTL3-dependent m6A modification is critical for the expression of key oncogenic proteins such as c-myc, SP1 and bcl-2 (Barbieri et al., 2017, Vu et al., 2017). In solid cancers, multiple studies show that m6A modifications are implicated in cancer progression and in acquired drug resistance (Zheng et al., 2019, Visvanathan et al., 2018). In addition, gene expression of the wider m6A writer complex, as well as other components of the m6A machinery, were shown to have a prognostic value, often in the context of anti-cancer immunity (Shen et al., 2021, Wu et al., 2021, Zhang et al., 2020).

Mammalian cells developed various mechanisms to detect and act against foreign RNA, which typically marks a viral infection. In order to recognise the presence of such RNA, the cells rely on common characteristic patterns such as double-stranded RNA (dsRNA) and the lack of a 5’-cap structure, rather than specific sequences. These patterns are detected by components of the innate immunity system, such as the cytosolic RNA sensors MDA5 and RIG-I and by Toll-Like Receptors (TLRs) located in the lysosome and on the plasma membrane. Upon activation, these receptors initiate signalling cascades that lead to production of interferon (IFN) and interferon-stimulated genes (ISGs) as well as activation of the NF-kB transcription factor and its target genes (Schlee & Hartmann, 2016). Importantly, endogenous RNA harbouring viral-like patterns can bind and activate the same RNA sensors and elicit similar signalling pathways (Hale, 2022). The origin of such endogenous RNA is often transposable element (TE) sequences. These TEs are embedded in the genome and are normally silenced, but can become transcriptionally active under certain circumstances, including epigenetic changes (Almeida et al., 2022).

We and others have previously shown that inhibition or loss of METTL3 leads to accumulation of dsRNA (Guirguis et al., 2023, Gao et al., 2020) and to activation of innate immunity pathways *in vitro* (Guirguis et al., 2023, Gao et al., 2020, Lu et al., 2020, Chen et al., 2019) and *in vivo* (Gao et al., 2020, Wang et al., 2020). Induction of type I IFN response has been shown to activate the adaptive immune system in multiple ways, including stimulating antigen presenting cell (APC) maturation and migration into lymph nodes, increasing CD8+ T-cell cytotoxicity and survival and shifting macrophages into a pro-inflammatory state (Zitvogel et al., 2015, Yu et al., 2022, Busselaar et al., 2024). Hence, innate immunity activation was found to enhance the activity of immune checkpoint inhibitors (ICI) (Heidegger et al., 2019, Ishizuka et al., 2019).

The interest in METTL3 as a drug target has led to the development of a number of small molecule inhibitors of METTL3 by Storm Therapeutics and others (Fiorentino et al., 2023, Wu et al., 2024, Li et al., 2025). The first of these inhibitors, STM2457, developed at Storm Therapeutics, was shown to be efficacious in models of AML, confirming the earlier findings using genetic depletion of METTL3, and validated METTL3 as a target for AML (Yankova et al., 2021). In addition, STM2457 and more recent METTL3 inhibitors such as STM3006 and STM3675 were shown to induce cancer cell-intrinsic formation of dsRNA and activation of type I IFN signalling *in vitro* and *in vivo* (Guirguis et al., 2023, Sturgess et al., 2023, Mayday et al., 2025), leading to anti-tumour immune responses and resulting in increased efficacy of anti-PD1 immunotherapy in preclinical models (Guirguis et al., 2023, Yu et al., 2024).

We recently described STC-15, a clinical stage, first in class, inhibitor of METTL3 (Patent number WO/2021/111124) that has been evaluated in a Phase 1 dose-escalation study in patients with advanced solid cancer malignancies (NCT05584111). STC-15, STM2457 and STM3675 molecules are derived from the same chemical series and share high similarity. In fact, STC-15 and STM3675 are identical except for a small change in the tail group and have similar potency. Hence, we used the two inhibitors interchangeably in our preclinical work and throughout this manuscript.

Here, we describe the molecular mechanism by which STC-15 and other inhibitors of this class induce dsRNA formation and activate a type I IFN response. We provide evidence that STC-15 drives strong anti-tumour immune responses and in combination with anti-PD1 induces profound cancer regression and durable immune memory.

## Results

### Loss of m6A from long exons following METTL3 inhibition results in intron retention and transcriptional run-on events

m6A modification levels were mapped transcriptome-wide in Caov3 ovarian cancer cells before and after METTL3 inhibition, using the quantitative and reproducible single-nucleotide resolution m6A-profiling GLORI approach (Fig. 1A, Fig. S1A-C) (Liu et al., 2023). After thresholding for sequencing coverage, we observed >70000 m6A sites with a pre-treatment modification level above 25% and noted a reduction in m6A modification at 99.4% of sites following STM3675 treatment, with over 92% of sites seeing a reduction to below 25% modification level (Fig. 1A-C).

**Figure 1.**
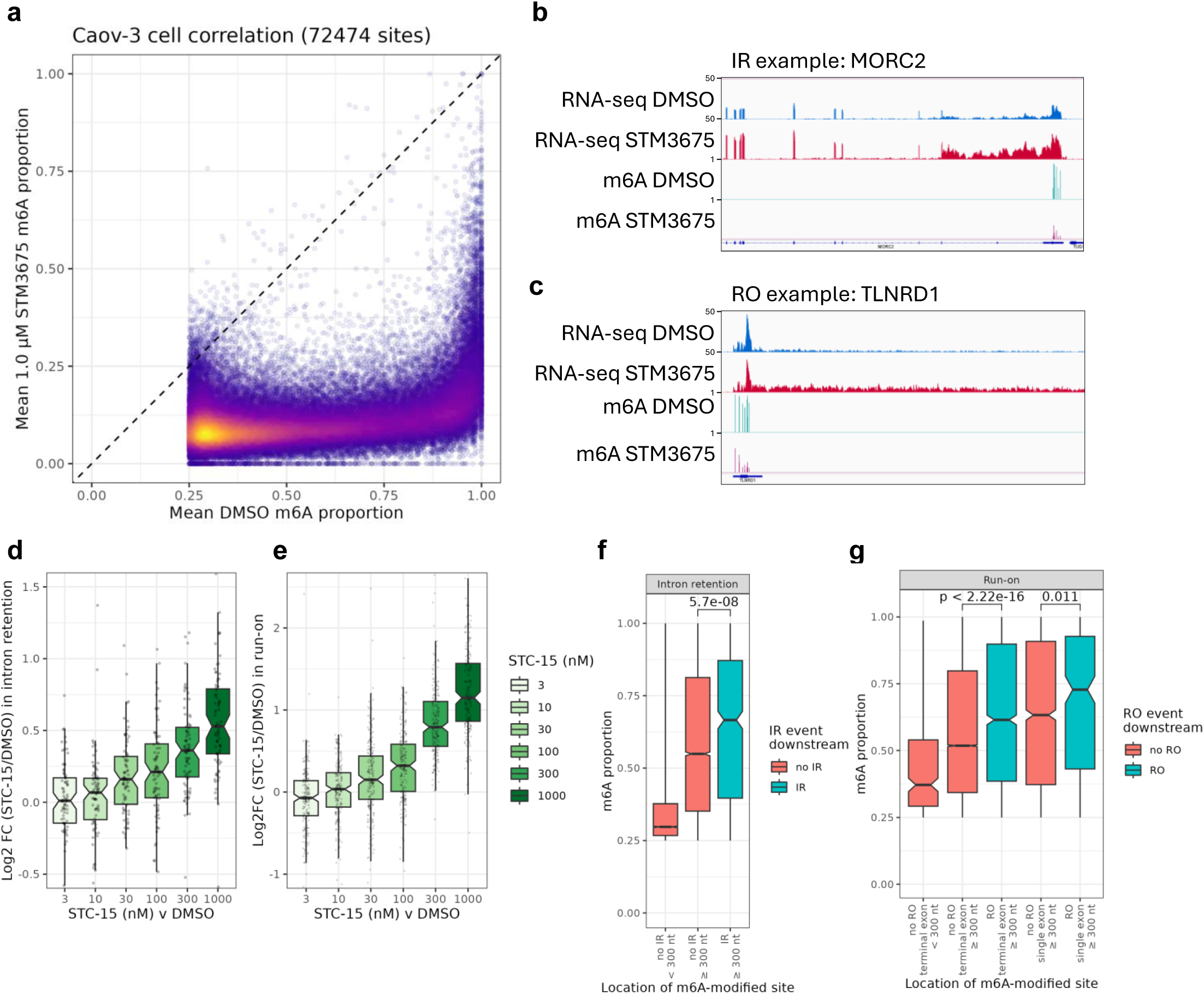
METTL3 inhibition leads to a dose-dependent increase in intron retention (IR) and run-on (RO) events downstream of m6A-enriched exons. **(a)** Scatter plot comparing the proportion of m6A modification at sites in Caov3 cells treated either with DMSO or 1 μM STM3675 for 24 h. Points are coloured from high (yellow) to low (purple) density. **(b-c)** IGV tracks showing RNA-seq and m6A proportions (GLORI data) from Caov3 cells treated for 24 h with either DMSO or 1 μM STM3675, exemplifying an IR event **(b)** and an RO event **(c)**. **(d-e)** Box and whisker plot of the log2 fold-change in IR **(d)** or RO **(e)** in Caov3 cells treated with 3, 10, 30, 100, 300 or 1000 nM STC-15 or 1 μM STM3675 relative to DMSO-treated cells at IR or RO significant events, respectively, as defined for the 1 μM STM3675 v DMSO comparison. **(f)** Box and whisker plots showing the distribution in the proportion of m6A modification at sites in short internal exons (< 300 nt), long internal exons not immediately upstream of significant IR events and long exons immediately upstream of significant IR events. **(g)** Box and whisker plots showing the distribution in the proportion of m6A modification at sites in short terminal exons, long terminal exons in multi-exon genes without significant RO events, long terminal exons in genes with significant RO events, single-exon genes without significant RO events and single-exon genes with significant RO events. Key pairwise comparisons between groups are indicated by brackets with corresponding p-values (Mann-Whitney U test).

We and others have previously shown that METTL3 depletion or its pharmacological inhibition in cancer cells results in the production of dsRNA structures and activation of a type I IFN response (Lu et al., 2020, Gao et al., 2020, Guirguis et al., 2023). As expected, treatment of Caov3 cells with STC-15 led to dose-dependent accumulation of dsRNA and induction of ISGs, as demonstrated by immunofluorescence assay utilising the dsRNA-specific J2 antibody (Weber et al., 2006) and an anti-IFIT1 antibody (Fig. S1D).

To identify potential sources of the accumulating dsRNA, we performed rRNA-depleted RNA-seq in Caov3 cells following METTL3 inhibition with STM3675. Analysis of the data revealed instances of aberrant transcript generation following treatment, including increased signal within a small subset of introns (245 introns) after controlling for increases in transcript expression, suggestive of disrupted splicing leading to intron retention (IR) events (Fig. 1B, Fig. S1E, Supplementary Table 1). In a different subset of genes (216 genes), we observed increased signal for thousands of bases downstream of the annotated 3’ UTR, suggestive of disrupted transcription termination leading to increased RNA polymerase II run-on (RO) events (Fig. 1C, Fig. S1F, Supplementary Table 1). In addition, upon METTL3 inhibition with a range of STC-15 concentrations, we detected a dose-dependent increase in the magnitude of these IR and RO events (Fig. 1D-E, Fig. S1G-H, Supplementary Table 1).

We noted that the IR events were enriched for events downstream of exons longer than 300 nt (79 events; 6.87 % of tested introns are downstream of long exons so the expectation would be ∼17), which have previously been shown to be m6A-enriched in other cell lines (He et al., 2023, Yang et al., 2022) (Supplementary Table 1). In our GLORI data, the m6A modification level at sites within internal and terminal exons longer than 300 nt and single-exon genes (all > 300 nt) was also greater than within shorter exons (Fig. 1F-G). We observed even higher m6A modification levels at m6A sites within long exons immediately preceding IR or RO events (Fig. 1F-G), suggesting that these misprocessing events are directly driven by m6A loss.

Pathway enrichment analysis of genes with either IR or RO events identified a small number of terms, including protein nuclear localisation in the IR gene set (Fig. S1I) and apoptotic signalling pathway genes in the RO set (Fig. S1J) (Supplementary Table 2).

### m6A levels differ across cell lines and correlate with the level of increase in IR/RO

To examine m6A profiles across cell types, we performed GLORI in five additional cell lines (Fig. 2A, Fig. S2A). The m6A profiles, as determined by GLORI, were strongly correlated between cell lines, with the more highly modified sites in one cell line also generally being highly modified in other cell lines. This is exemplified by comparison between Caov3 and A549 m6A-site modification levels (Fig. 2B). However, a global difference in the m6A modification level was found between cell lines, with most sites showing higher modification in Caov3 compared to A549 (Fig. 2A-B).

**Figure 2.**
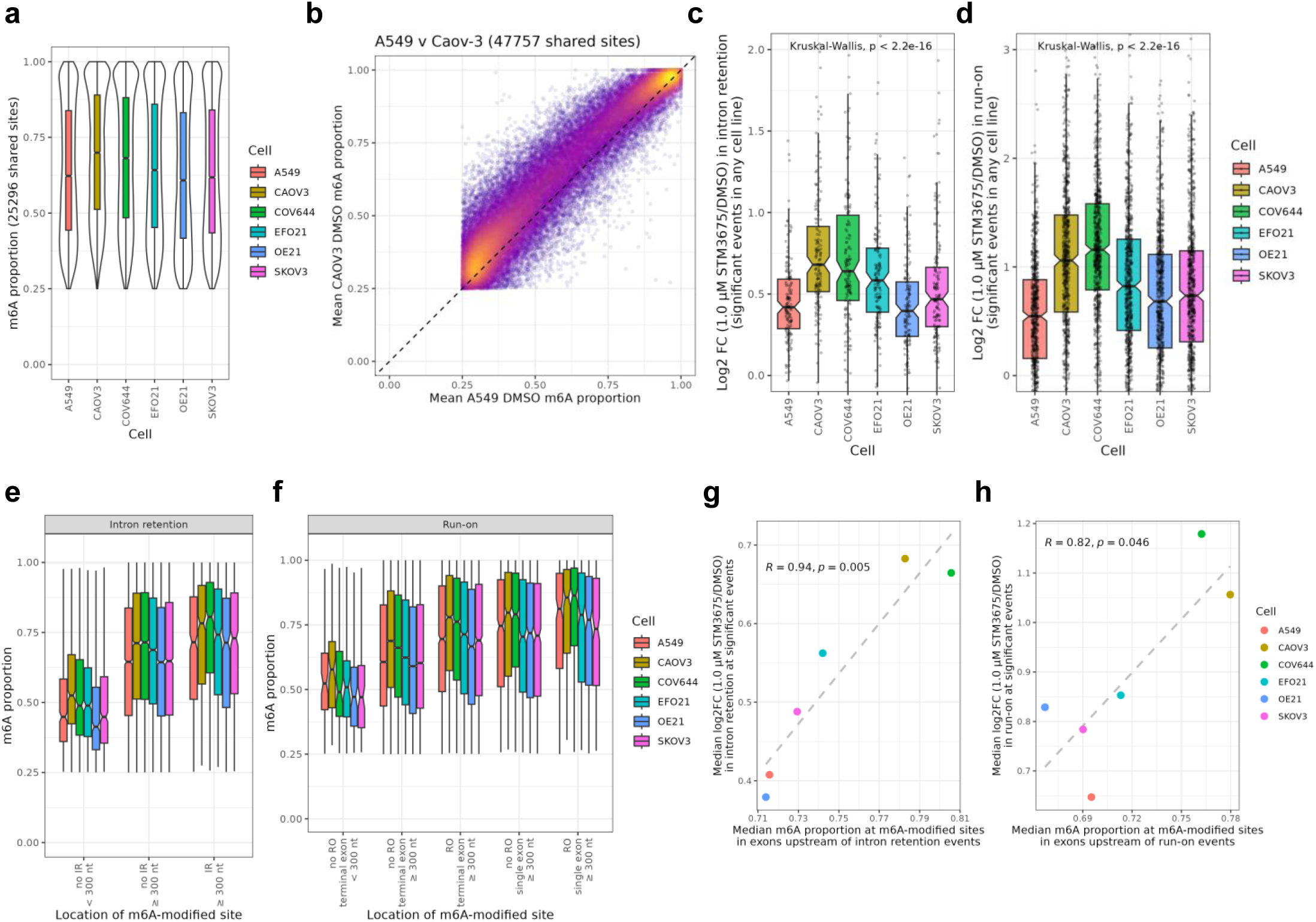
The level of m6A in the upstream exon correlates with the level of IR and RO events across cell lines. **(a)** Violin and box and whisker plots showing the distribution of m6A proportions in a range of DMSO-treated cell lines for sites passing thresholds (m6A proportion ≥ 0.25; combined unmodified and modified A count ≥ 20) in all cell lines. **(b)** Scatter plot comparing the proportion of m6A modification in DMSO-treated Caov3 and A549 cells for sites passing thresholds (m6A proportion ≥ 0.25; combined unmodified and modified A count ≥ 20) in both cell lines. Points are coloured from high (yellow) to low (purple) density. **(c-d)** Box and whisker plots of the log2 fold-change in IR **(c)** or RO **(d)** in 6 cell lines treated with 1 μM STM3675 vs DMSO-treated cells at IR or RO significant events in any cell line. **(e)** Box and whisker plots showing the distribution in the proportion of m6A modification for each cell line at sites in short internal exons (< 300 nt), long internal exons not immediately upstream of significant IR events and long exons immediately upstream of significant IR events. **(f)** Box and whisker plots showing the distribution in the proportion of m6A modification for each cell line at sites in short terminal exons, long terminal exons in multi-exon genes without significant RO events, long terminal exons in genes with significant RO events, single-exon genes without significant RO events and single-exon genes with significant RO events. **(g)** Scatter plot showing the relationship across cell lines between the median log2 fold-change in IR at significant events with m6A sites in the upstream exon and the median m6A proportion of these m6A sites. **(h)** Scatter plot showing the relationship across cell lines between median log2 fold-change in RO at significant events with m6A sites in the upstream exon and the median m6A proportion of these m6A sites.

We next identified IR and RO events in RNA-seq data from these cell lines and again observed a difference in the magnitude of these events across cell lines (Fig. 2C-D, Fig. S2B-D, Supplementary Table 3). The median m6A levels in long exons immediately preceding these IR and RO events were higher than in other long exons in all cell lines, with the trend in different m6A modification levels across different cell lines following the trend observed for global m6A levels (Fig. 2A, E, F). Remarkably, we observe a significant correlation between the median level of m6A in exons upstream of either IR or RO events and the median fold change in signal for these events (Fig. 2G-H). This provides further support that m6A loss may directly drive these misprocessing events and suggests that baseline m6A levels can predict the level of these IR/RO events upon METTL3 inhibition.

### METTL3i-induced IR/RO events generate transcripts forming double stranded RNA

The IR and RO events produce transcripts containing intronic or intergenic regions that are often rich in TEs and other repetitive sequences (Wells & Feschotte, 2020). These regions would not normally be included in mature mRNA and may result in formation of dsRNA, leading us to speculate that these aberrant transcripts might be the source of the dsRNA arising from METTL3 inhibition. To test this hypothesis, we characterised METTL3i-induced dsRNA transcripts in an unbiased manner. We performed dsRNA immunoprecipitation and sequencing (dsRIP-seq) experiments using the J2 antibody (Gao et al., 2021, Coban et al., 2024) (Fig. 3A) on RNA purified from Caov3 cells treated for 24 hours with 1 μM STM3675 or DMSO control. To validate our approach, we compared the dsRIP-seq samples to matched RNA-seq input in the control DMSO condition and found significant enrichment of known dsRNA transcripts, including BPNT1 and BRI3BP (Sakurai et al., 2014, Ahmad et al., 2018), but a depletion of transcripts from several housekeeping genes (Fig. S3A-B).

**Figure 3.**
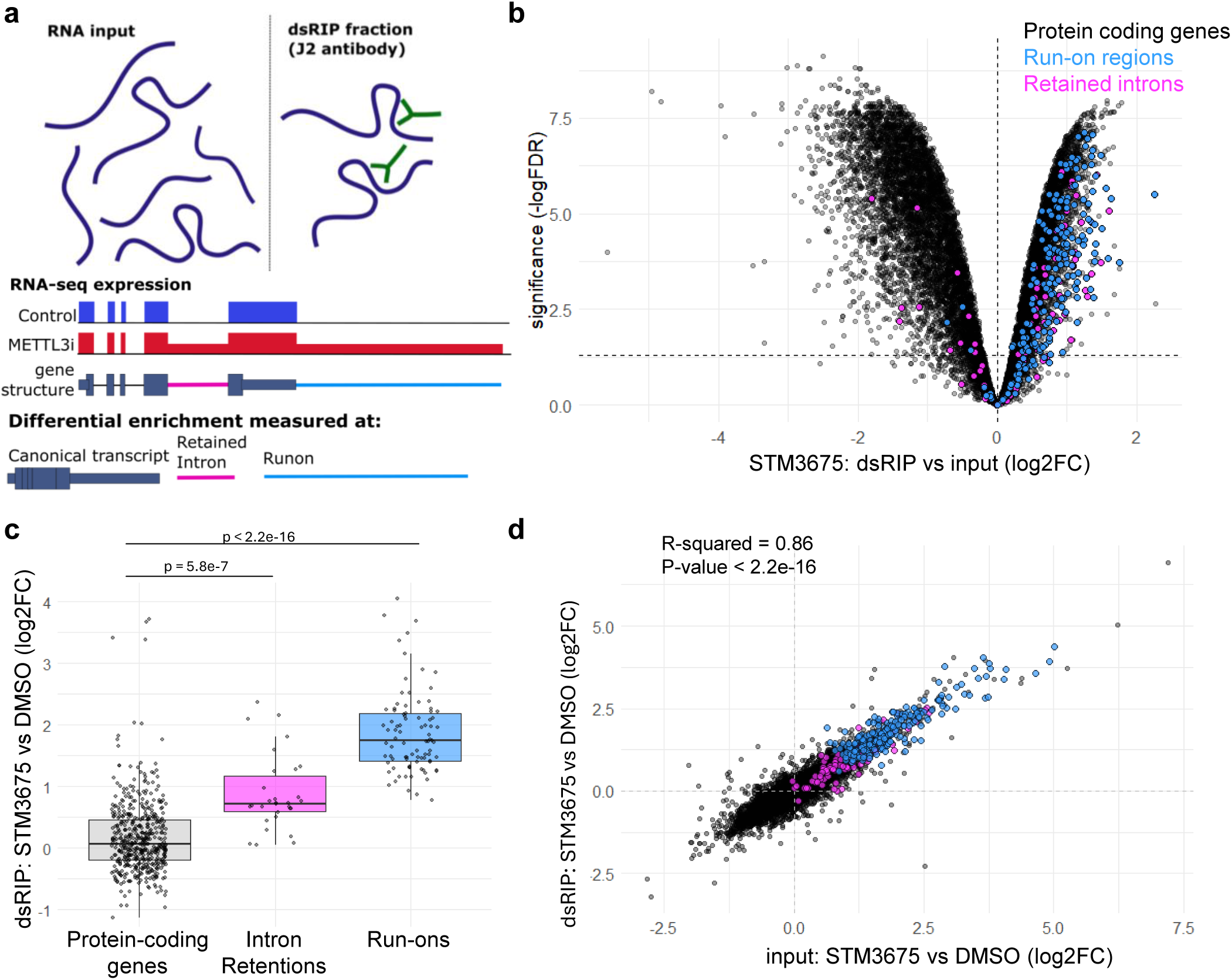
IR/RO events produce dsRNA-rich aberrant transcripts. **(a)** Diagram illustrating dsRIP-seq. Doublestranded RNA is immunoprecipitated and then sequenced alongside input RNA. Analysis was performed on the canonical transcript, as well as on the previously identified retained intron (IR) and run-on (RO) regions independently. **(b)** Enrichment of RNA species in the J2 dsRIP-seq compared to input, on Caov3 cells treated for 24 h with 1 μM STM3675. Canonical transcripts are shown in grey, IRs in pink, and ROs in blue. **(c)** Box and whisker plot of the fold-change in dsRIP pulldown after 24 h treatment with 1 μM STM3675 vs DMSO. Only species that are classed as dsRNA in either DMSO or STM3675 are included (dsRIP vs input: log2FC > 1, FDR < 0.05). Canonical protein coding transcripts, IRs and ROs are displayed separately. **(d)** Differential expression in input is plotted against differential expression in dsRIP after treatment for 24 h with 1 μM STM3675 or DMSO. IRs in pink, and ROs in blue.

To examine the enrichment of the IR/RO events, we generated a genome annotation to handle each of these regions as if it were an independent transcript for differential expression analysis alongside the canonical gene transcripts (Fig. 3A) (see methods). Examining the STM3675-treated samples, we found that many IR and RO regions were significantly enriched by dsRIP compared to input (Fig. 3B, Supplementary Table 4), suggesting that these regions contain dsRNA structures. Indeed, comparing the dsRIP-seq for STM3675-treated vs DMSO-treated samples, we found that the IR and RO regions accounted for the majority of the most increased RNA species (Fig. 3C).

An increase in dsRIP capture of a given transcript following METTL3 inhibition can be explained in two ways. If m6A modification prevents complementary sequences from forming dsRNA (Spitale et al., 2015), m6A removal will result in an increase in ‘double-strandedness’, i.e. the propensity of a given transcript to form dsRNA structures. Alternatively, the increase in dsRIP capture may reflect higher expression level of a dsRNA transcript that is already double-stranded. To distinguish between these possibilities in STM3675-versus DMSO-treated samples, we compared the change in dsRIP enrichment for each transcript with the corresponding change in input RNA level (Fig. 3D, Fig. S4A, Supplementary Table 4). Very few transcripts showed any change in double-strandedness, arguing against the idea that m6A generally inhibits the formation of dsRNA by blocking complementary sequence hybridization. The IR and RO regions themselves also typically did not show a change in double-strandedness. Nevertheless, several of the transcripts with increased double-strandedness corresponded to canonical transcripts of genes that undergo IR or RO events (Fig. S4B). This supports a model in which the IR and RO regions are physically connected to the canonical transcript, such that dsRNA formation within these regions leads to J2 antibody enrichment of the entire transcript. The magnitude of this effect is likely to scale with the fraction of transcripts that contain the misprocessing event, since a higher proportion of misprocessed isoforms would increase the overall likelihood of capturing the transcript in dsRIP. Notably, some other genes with changes in double-strandedness are accounted for by alternative splicing (WTAP, Wei et al., 2021) or alternative polyadenylation events (SDR42E1) upon METTL3i (Fig. S4C).

We found that change in RNA expression level accounts for the majority of changes in dsRIP capture (Fig. 3D). The IR and RO events are highly upregulated upon STM3675 treatment compared to the low baseline expression in the DMSO-treated sample, and therefore contribute the biggest increase in dsRIP-captured species. Overall, our findings suggest that rather than a change in transcript double-strandedness, the production of new transcripts carrying IR/RO regions is the primary source of increased dsRNA upon METTL3 inhibition.

### Inverted Alu repeats in IR and RO regions are bound by the dsRNA sensor MDA5 in the cytoplasm

While our dsRIP approach identified transcripts with dsRNA structures, only those dsRNAs that reach the cytoplasm are likely to induce an innate immune response, so we repeated the dsRIP-seq experiment using a cytoplasmic fraction as the input. Furthermore, we did not purify the RNA before the immunoprecipitation, to ensure that we only enriched for species that form dsRNA in their endogenous cellular state. We confirmed that in the cytoplasm, many IR and RO events are enriched by dsRIP and these contribute most of the largest increases in the dsRNA pool (Fig. S5A-B, Supplementary Table 5).

Since not all IR and RO events were enriched by dsRIP-seq, we decided to examine their distinguishing features. The IR/RO regions frequently include TEs, so we examined the number and identity of TEs in the double-stranded and non-double stranded IR/ROs. We found that dsRIP-enriched IR/RO regions contained significantly more examples of inverted Alu repeats than IR/RO regions that were not enriched by dsRIP (Fig. 4A, Supplementary Table 6). Alu elements are a family of Short Interspersed Nuclear Element (SINE), themselves a class of TE, that, when found in the formation of inverted pairs, have been reported to fold back on themselves to form dsRNA structures (Ahmad et al., 2018, Athanasiadis et al 2004, Kawahara & Nishikura, 2006).

**Figure 4.**
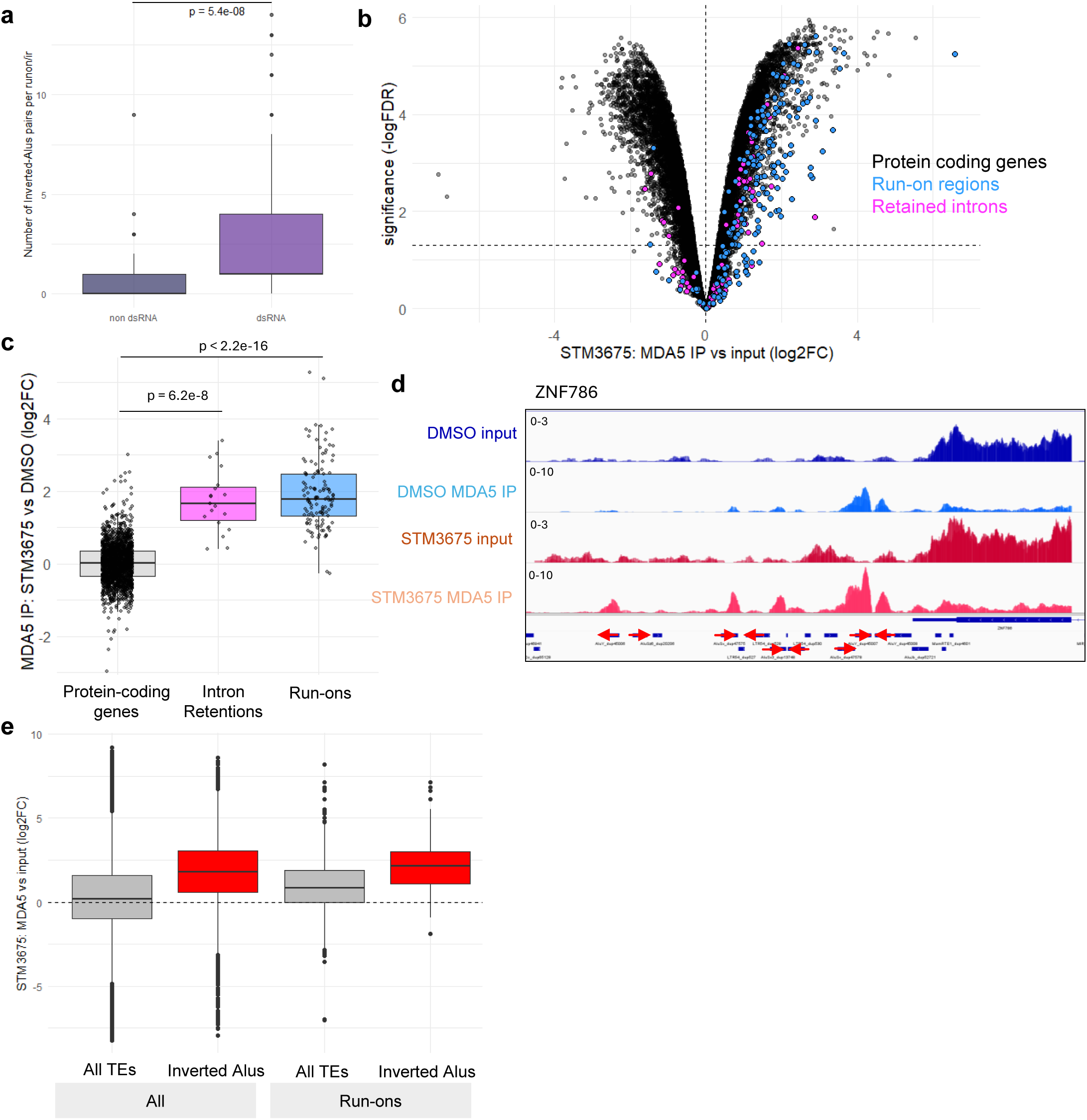
RO regions are enriched in MDA5 IP upon STM3675 treatment, with binding at inverted Alu pairs. **(a)** Number of inverted Alu pairs per IR/RO region, divided into those that do not form dsRNA (not enriched in the J2 dsRIP) and those that do form dsRNA (enriched in the J2 IP). The dsRNA IR/Ros contain significantly more inverted Alus sequences than non-pulled dsRNA IR/Ros. **(b)** Enrichment of RNA species in the MDA5 RIP experiment compared to input, from the cytoplasmic fraction of Caov3 treated for 24 h with 1 μM STM3675. Canonical transcripts are shown in grey, IRs in pink, and ROs in blue. **(c)** Box and whisker plot of the foldchange in MDA5 RIP pulldown after 24 h treatment with 1 μM STM3675 vs DMSO. Only species that are classed as bound by MDA5 in either DMSO or STM3675 are included (MDA5 RIP vs input: log2FC > 1, FDR < 0.05). Canonical protein coding transcripts, IRs and ROs are displayed separately. **(d)** Input RNA-seq and MDA5 RIP-seq IGV tracks for example gene ZNF786 in DMSO and STM3675-treated Caov3 cells. Digestion with RNase T1 results in enrichment at dsRNA regions, overlapping with inverted Alu repeats. **(e)** Enrichment of MDA5 RIP-seq over input RNA-seq after STM3675 treatment, for all TEs and for inverted Alus.

One of the key cytoplasmic dsRNA sensors is MDA5, which has previously been reported to recognise dsRNA formed by endogenous inverted Alu repeats (Ahmad et al., 2018). Hence, we examined whether the RO and IR events are bound by MDA5 in the cytoplasm, using MDA5 RIP-seq (Fig. S6A). MDA5 is reported to bind promiscuously to single-stranded RNA (ssRNA), but only forms filaments upon cooperative binding to dsRNA (Ahmad et al., 2018). We performed MDA5 RIP on Caov3 cells treated with 1 μM STM3675 or DMSO control, then used RNase T1 to digest the ssRNA, enriching for dsRNA regions. We examined the 3’ UTRs of transcripts known to be bound by MDA5 (Ahmad et al., 2018), and found MDA5 IP enrichment compared to input RNA-seq at inverted Alu repeats (Fig. S6B).

To explore the changes in MDA5-bound transcripts in STM3675- vs DMSO- treated Caov3 cells, we first examined MDA5 RIP-seq enrichment of each transcript or IR/RO event compared to cytoplasmic input RNA-seq (Fig. 4B). We found that many ROs were bound by MDA5, and they contributed most of the changes in MDA5 binding upon METTL3 inhibition (Fig. 4C). However, the RNase T1 digestion degrades the ssRNA regions of each transcript, which makes the overall enrichment of the transcript dependent on the number of dsRNA regions (Fig. 4D). Therefore, we examined MDA5 enrichment at each TE within an IR or RO event and found that inverted Alu pairs in these regions were highly enriched in the MDA5 RIP-seq (Fig. 4E, Fig. S6C, Supplementary Table 7). This suggests that the inverted Alus in the IR and RO regions are bound by MDA5, supporting our finding that these regions are the source of dsRNA induced by METTL3 inhibition.

### METTL3 inhibition induces a profound type I interferon response that correlates with the extent of m6A methylation and the occurrence of IR/RO events

Pathway enrichment analysis of differentially expressed genes in Caov3 cells following METTL3 inhibition revealed a dose-dependent induction of a transcriptional programme reflecting a cell-intrinsic innate immune response, such as pathways related to IFN, NF-kB signalling and viral response (Fig. 5A, Fig. S7A-B, Supplementary Table 8). These pathways were also induced in the majority of the 6 cell lines tested (Fig. 5B, Fig. S7C-D, Supplementary Table 8), as exemplified by a heatmap of RNA level changes of transcripts within the Negative Regulation of Viral Process GO term (Fig. 5B). The differences in the induction level of genes associated with this pathway broadly reflects differences in the level of IR/RO induction upon METTL3 inhibition and differences in baseline m6A modification levels across these cell lines. For example, Caov3 and COV644 have higher m6A modification levels, higher IR/RO and stronger viral response pathway induction than A549 (Fig. 2, Fig. 5B, Fig. S7C-D). The exception being OE21 with comparable viral response pathway induction to Caov3 and COV644 but with IR/RO induction and baseline m6A modification levels similar to A549. However, OE21 cells already show high ISG transcription levels at baseline (ISG^High^ cell line, Liu et al., 2018), which may explain their higher sensitivity to further IFN induction.

**Figure 5.**
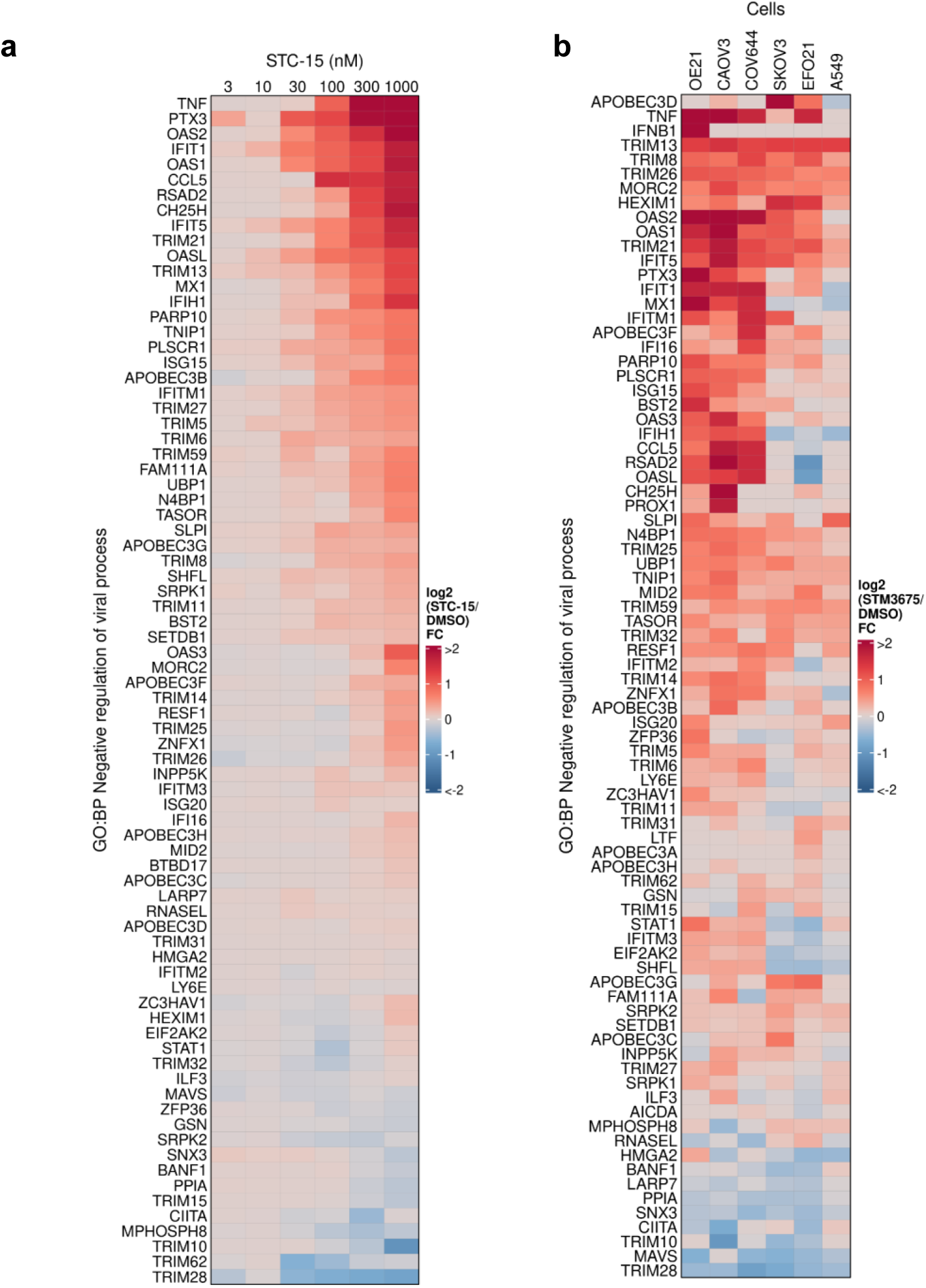
METTL3 inhibition leads to increased expression of innate immune genes. **(a)** Heatmap showing the log2 fold-change in RNA level of genes within the biological process GO term negative regulation of viral process in Caov3 cells treated with 3, 10, 30, 100, 300 and 1000 nM STC-15 vs DMSO-treated cells. **(b)** Heatmap showing the log2 fold-change in RNA level of genes within the biological process GO term negative regulation of viral process in 6 cell lines treated with 1 μM STM3675 vs DMSO-treated cells.

We validated the induction of IFN signalling at the RNA and protein level in Caov3 cells over time following treatment with STC-15. We observed a gradual increase in IFNβ transcript level, peaking at 48h post METTL3 inhibition (Fig. S8A) and a strong induction of ISG transcription (Fig. S8B). A concomitant upregulation of ISG on the protein level was observed, as well as the phosphorylation of IRF3 and STAT1, both acting upstream of ISG induction in the IFN signal transduction cascade (Fig. S8C). In line with these observations, we found a time-dependent increase in secreted IFNβ and CXCL10 (Fig. S8D). Co-treatment with STC-15 and Ruxolitinib, a JAK1/2 inhibitor, or BAY-985, a TBK1/IKKε inhibitor, blocked STAT1 phosphorylation and downstream signalling (Fig. S8E and Fig. S8F, respectively), further demonstrating the activation of a classical dsRNA-induced IFN response.

### STC-15 induces T-cell mediated anti-tumour responses in an *in vitro* model of immune-cell mediated tumour cell killing

To investigate whether induction of a dsRNA-driven IFN response could activate immune-cell mediated anti-tumour responses, we employed a co-culture system of fluorescently labelled SKOV3 cells, cultured in the absence or presence of donor PBMC. Cultured in isolation, SKOV3 cell growth and viability was only modestly reduced by METTL3 inhibition via STC-15 (Fig.6A, left panels). However, when co-cultured with PBMC, STC-15 treatment induced a clear dose-dependent decrease in SKOV3 cell numbers and increased apoptosis, leading to complete elimination of SKOV3 cells at 72 h post treatment (Fig. 6A, right panels). Importantly, PBMC-mediated SKOV3 killing was induced at very low STC-15 concentrations and exceeded the effect of the positive control (high dose IL-2), suggesting that anti-tumour immune activation is exquisitely sensitive to METTL3 inhibition. We also detected a dose-dependent increase in the secretion of IFNg, TNFa and IL-1b proinflammatory cytokines in the co-culture system (Fig. S9A).

**Figure 6.**
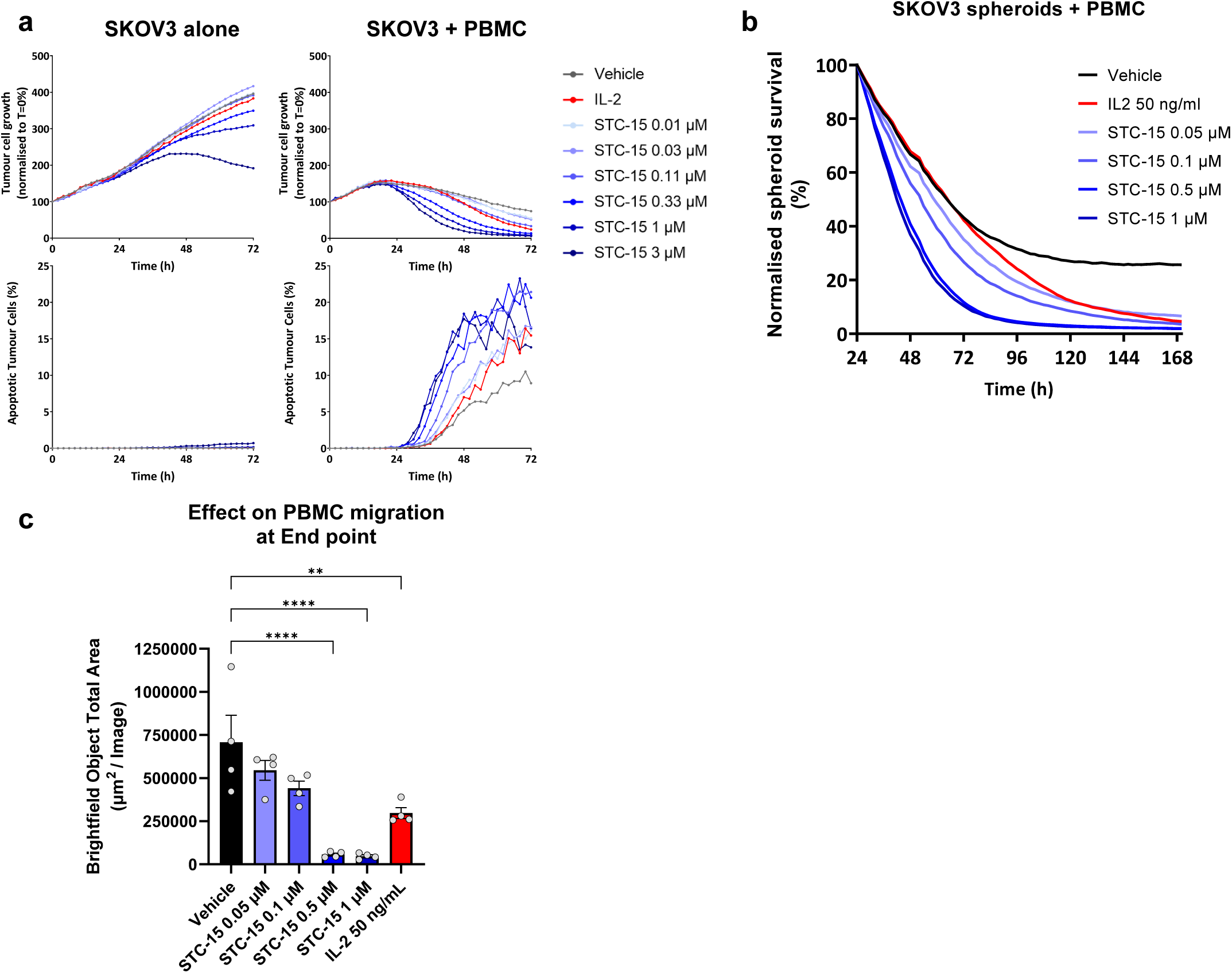
STC-15 induces PBMC-mediated killing of cancer cells in a co-culture system. **(a)** SKOV3 co-culture system. Top: Tumour cell growth of SKOV3-NLR cell line in presence of DMSO or varied doses of STC-15 in the absence (left) and presence (right) of donor PBMCs. Bottom: induction of apoptosis in SKOV3 cells under the same conditions. Data are normalised to T=0. Data from one donor is presented and is representative of all three donors. **(b)** SKOV3-NLR cells were seeded in ultra-low attachment plates to generate 3D spheroids. After 48h, donor PBMC and STC-15 or control treatment were added and incubated for additional 7 days. Spheroid size was monitored by the IncuCyte every 2 h. Data are normalised to T=0. Data from one technical replicate of one donor is presented and is representative of all three donors. **(c)** Bar plot is showing the quantification of PBMC migration into the centre of the well. Data are presented as mean ± SEM. Each dot represents a technical replicate. Data from one donor is presented and is representative of all three donors tested. Statistical analysis was performed using ANOVA testing with multiple comparisons. Each group was compared to Vehicle control. Statistically significant comparison are denoted by asterisks (**, p<0.005; ****, p<0.0001).

In a 3-Dimensional variation of the co-culture system, SKOV3 cells were grown as spheroids. A clear dose-dependent effect on spheroid size was observed, even at low STC-15 concentrations (Fig. 6B, Supplementary Video 1-2). Examining the co-culture images (Fig. S9B) revealed a strong dose-dependent effect of STC-15 treatment on PBMC migration toward the spheroid body (Fig. 6C, Fig. S9C).

Importantly, we could not detect a significant effect on cell viability or T-cell proliferation when PBMC cultured in isolation were treated with STC-15 and stimulated with CD3/CD28 (Fig. S9D), in line with previous observations (Guirguis et al., 2023). Notably, the killing of cancer cells by STC-15 in the co-culture system was mediated by immune cells, as SKOV3 cell viability was not affected by STC-15 when grown alone. Thus, our co-culture experiments suggest that STC-15 effect on immune activation is governed by a cancer-intrinsic mechanism, which then enhances the PBMC-mediated killing of the cancer cells.

### STC-15 monotherapy inhibits tumour growth *in vivo*, and induces tumour regression in combination with anti-PD1 checkpoint therapy

To test STC-15 activity *in vivo*, we used two mouse syngeneic models that can capture immune-related mechanisms of action: an A20 lymphoma model (Fig. S10) and an MC38 colorectal cancer model (Fig. 7, Fig. S11) In the MC38 model, high-dose STC-15 monotherapy treatment showed a significant effect on tumour growth, comparable to anti-PD1 therapy, while low-dose STC-15 only had a minor effect (Fig. 7A-B). STC-15 *in vivo* efficacy was dependent on the presence of the immune system, as we observed a complete lack of response in immune-compromised NRG compared to immune competent C57BL/6 mice (Fig. 7C). In addition, depletion of CD8+ cytotoxic T-cells reversed STC-15 efficacy (Fig. S10A-B).

**Figure 7.**
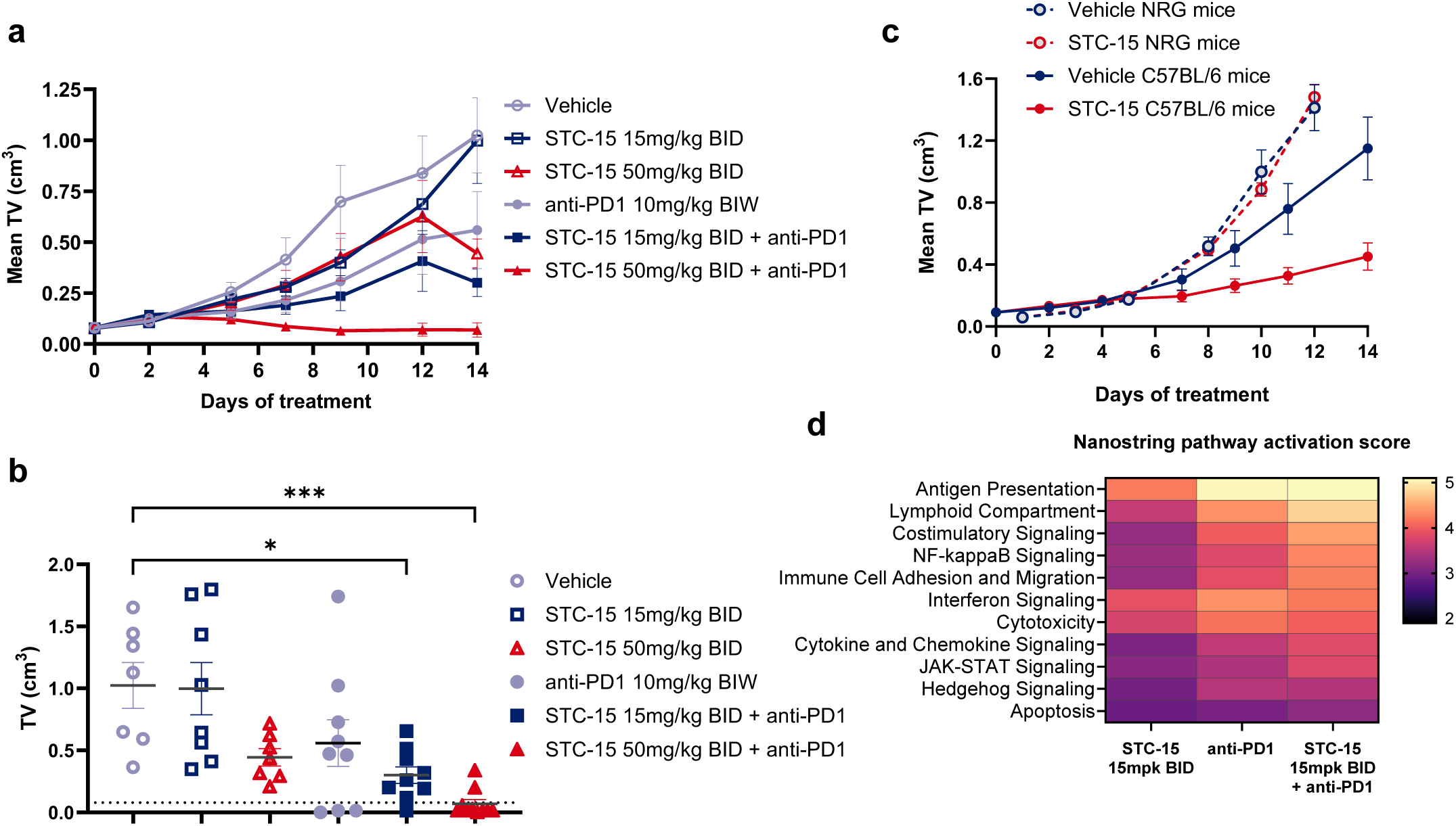
STC-15 efficacy in syngeneic mouse models. **(a-b)** Mean tumour volume over time **(a)** and individual tumour volume at End Point **(b)** of MC38 sub-cutaneous tumours implanted immune competent host (C57BL/6 background). Mice (n= 10 / group) were treated with the indicated treatments and regimens. Dotted line denotes average tumour volume at baseline. Data are presented as mean ± SEM. Statistical analysis was performed using ANOVA testing with multiple comparisons. Each group was compared to Vehicle control. Statistically significant comparison are denoted by asterisks (*, p<0.05; ***, p<0.0005). **(c)** Volume of MC38 sub-cutaneous tumours implanted in immune compromised host (NRG background) or immune competent host (C57BL/6 background). Mice (n= 10 / group) were treated with either vehicle control or 100 mg/kg STC-15 QD. Data are presented as mean ± SEM. **(d)** From the same experiment, tumours (n=3) from the indicated groups were resected at End Point and gene expression was assessed using the Nanostring IO-360 panel. Top pathway activation scores, as calculated by nSolver software, are presented.

We next investigated whether STC-15 could induce similar IR and RO events *in vivo*. The MC38 mouse syngeneic colorectal cancer model was implanted subcutaneously and the mice were dosed once daily with either vehicle or 100 mg/kg STC-15. Whole blood and tumour samples were collected at several time points after 7 days of daily dosing, and RNA-seq was performed. We observed IR and RO events in both sample types, suggesting a conservation of the mechanism in mouse cells *in vivo*. The magnitude of these IR/RO events was typically greater and longer lasting in tumour comparing to blood samples (Fig. 8A-C, Supplementary Table 9). In addition, we noted increased level of transcripts that are part of the viral response pathway for up to 120 hours after the last dose, particularly in tumour samples (Fig. 8D, Supplementary Table 8).

**Figure 8.**
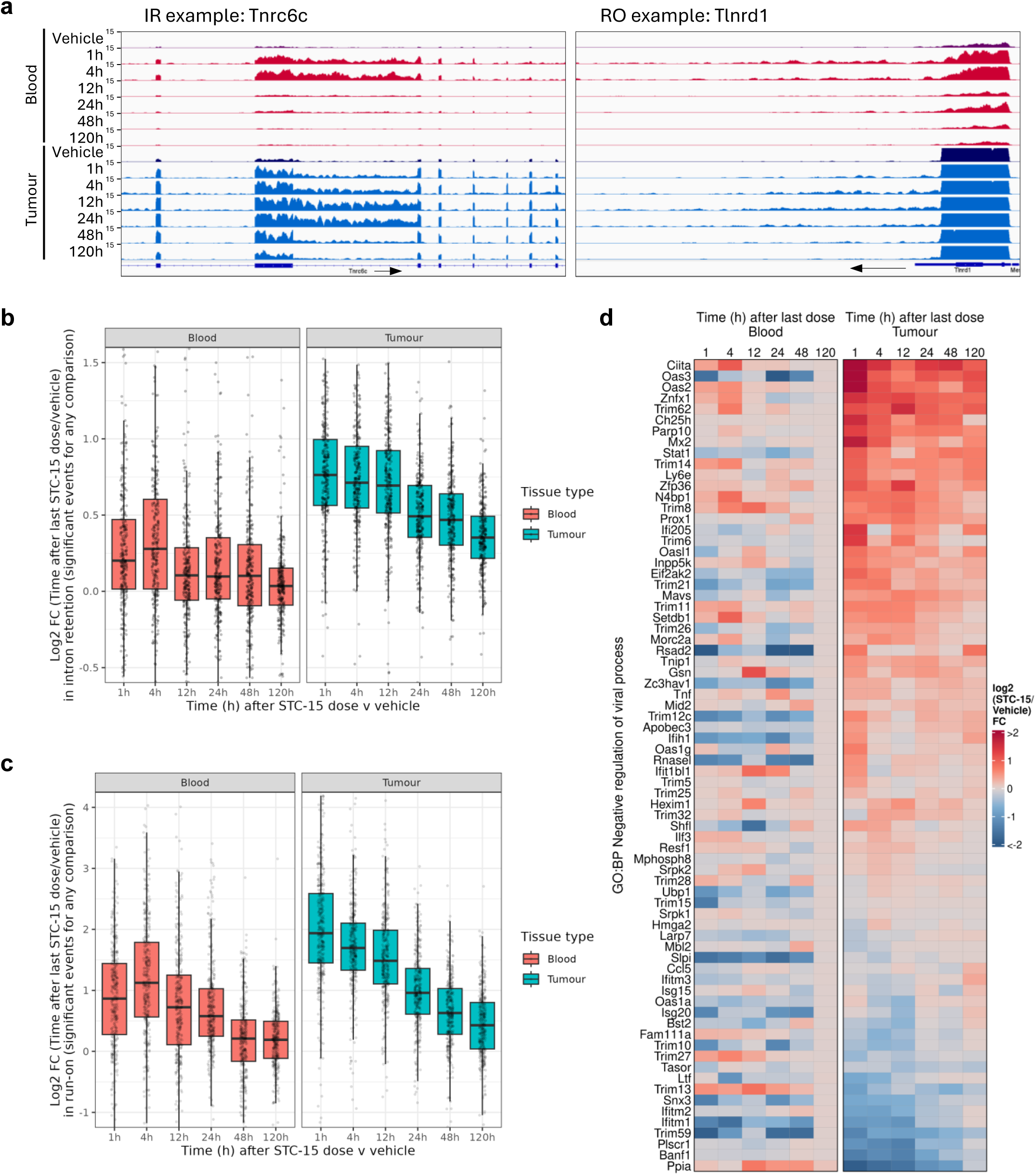
STC-15 leads to increased IR/RO and increased innate immune transcript levels in in vivo samples. **(a)** IGV tracks showing RNA-seq from whole blood or MC38 tumour cells from C57BL/6 mice treated with vehicle or 100 mg/kg STC-15 at 1, 4, 12, 24, 48 or 120 hours after the last dose, exemplifying an IR event (leftpanel) and an RO events (right panel). **(b-c)** Box and whisker plots of the log2 fold-change in IR **(b)** or RO **(c)** in whole blood or MC38 tumour cells treated with 100 mg/kg STC-15 at time points after the last dose v vehicle-treated mice. **(d)** Heatmap showing the log2 fold-change in RNA level of genes within the biological process GO term negative regulation of viral process in whole blood or MC38 tumour cells treated with 100 mg/kg STC-15 at time points after the last dose vs vehicle-treated mice.

We hypothesised that STC-15 efficacy could be further enhanced by combining STC-15’s innate immune activating effect with T-cell activation by a checkpoint inhibitor. We therefore combined the two treatments, co-dosing STC-15 with anti-PD1. Indeed, the combination of low-dose STC-15 and anti-PD1 treatment resulted in significantly improved tumour growth inhibition, while the combination of high-level STC-15 and anti-PD1 therapy led to complete tumour regression in the majority of mice (Fig. 7A-B, Fig. S11A). As these tumours were too small to resect, we focussed our analysis on the remaining tumours from the groups treated with either vehicle, low dose STC-15, anti-PD1, or the combination of low dose STC-15 and anti-PD1. A comparative gene expression analysis was performed, using the Nanostring IO-360 panel, with the top enriched pathways presented in Fig. 7D. The low-dose STC-15 group showed modest enrichment pathway scores, in line with the minimal effect on tumour volume in this group. The expression of the same pathways was highly enhanced in tumours from the combination arm, compared to either STC-15 or anti-PD1 monotherapy. One of the most upregulated genes in all three treatment groups was *CD8a* (Fig. S11B), suggesting increased CD8^+^ T cell infiltration in the tumours. Indeed, this was confirmed by CD8^+^ immunohistochemistry (IHC) analysis of tumour samples (Fig. S11C).

## Discussion

STC-15 and its research analogue STM3675 are potent and selective METTL3 inhibitors, with STC-15 being the first inhibitor of an RNA modifying enzyme to enter human clinical development. In this manuscript, we quantitate the level of m6A across cell lines and characterise the effects of METTL3 inhibition on these levels at single site resolution. We describe the resulting RNA misprocessing, leading to intron retention and transcriptional run-on events downstream of a subset of exons that are particularly highly m6A-modified. We use immunoprecipitation experiments to pinpoint these events as the source of the dsRNA previously reported to be induced by METTL3 inhibition and show that they are bound by the cytoplasmic dsRNA sensor MDA5. We characterise the downstream innate immune signalling induction and show that these effects can be harnessed to activate the adaptive immune system and drive anti-tumour immunity *in vitro* and *in vivo*, providing a rationale to combine STC-15 and ICI in the clinic.

We identify IR and RO events that occur after METTL3 inhibition specifically downstream of a subset of long, highly m6A-modified exons. Although the exact molecular mechanism driving these events remains unknown, it is likely associated with changes in splice site recognition and 3’ end processing. m6A loss has previously been linked to alternative splicing events after METTL3 depletion in mouse embryonic stem cells (Wei et al., 2021), as well as in lung cancer cell lines, where intron retention events occur upon depletion of the m6A reader YTHDC1 (Elvira-Blazquez et al., 2024). m6A also affects 3’ end processing: it has recently been shown that the loss of VIRMA, a component of the m6A writer complex, leads to 3’ UTR lengthening (Yue et al. 2018), while inhibition of METTL3 results in a switch to longer 3’ UTRs in bladder cancer (Koch et al., 2025). The RO events described here presumably occur due to skipping of proximal polyadenylation sites, and result in transcription of a much longer region. They are reminiscent of the Downstream of Gene (DoG) transcripts that are induced by osmotic stress, heat shock and viral infection (Vilborg et al., 2015, Vilborg & Steitz, 2017, Morgan et al., 2022), however it remains unclear whether DoGs can be triggered by additional signals. Notably, we found RO events in a different specific subset of genes, which commonly have higher m6A modification levels at m6A sites within long exons immediately preceding the RO event. While the molecular mechanisms behind the changes in splice site and polyadenylation site usage remain unknown, the events appear to be linked to gene structure rather than function. In our dataset, homologous genes with a similar structure often experience the same IR or RO events and some are conserved across species (e.g. TLNRD1).

In this study, we have focused on the role of IR and RO events in forming dsRNA, but these aberrant transcripts may also result in defects in protein production from the originator gene. A retained intron might result in nuclear retention, a premature stop codon, or a frame shift, causing a reduction in expression of the full length protein (Elivra-Blazquez et al., 2024).

Moreover, improper 3’ end processing and polyadenylation may result in changes in regulatory sequences affecting RNA stability. Unlike DoGs, which have been shown to be retained in the nucleus (Vilborg et al., 2015, Vilborg & Steitz, 2017), we find that transcripts containing ROs are present in the cytoplasm and bind the cytoplasmic dsRNA sensor MDA5, although some reduction in nuclear export may occur. These aberrant transcripts, if translated, may also contribute to neoantigen generation.

dsRNA from exogenous sources such as viruses is bound by cytoplasmic RNA sensors like MDA5 and induces an innate immune response (Schlee & Hartmann, 2016). We show that both IR and RO events frequently include pairs of inverted Alu repeats, that fold to form stretches of ∼300 bp dsRNA, and are bound by MDA5. Upregulation of endogenous inverted Alu repeats via other pathways such as EZH2 inhibition (Feng et al., 2024) and DNA demethylating therapies (Mehdipour et el., 2020, Hosseini et al., 2024), have also been shown to activate innate immune responses, which could be rescued by disabling the dsRNA sensing pathways (Feng et al., 2024, Hosseini et al., 2024). We focused on MDA5 due to its known binding affinity for inverted Alu pairs, but other dsRNA sensors such as PKR and RIG-I may also play a role.

Inverted Alu pairs are found in the 3’ UTRs of some mRNAs under normal conditions, so the cell needs mechanisms to prevent the deleterious effects to cells and tissues. Our RNA-seq data show that many of the IR/RO regions are already expressed, albeit at a very low level, at baseline conditions, and upon METTL3i treatment their expression is upregulated. This suggests that the triggering of the innate immune response is quantitative rather than qualitative, and a certain threshold needs to be crossed in order to effectively trigger this response. The threshold may be different in different cancer models, and is likely to be related to the baseline expression level of RNA sensors, such as MDA5. Indeed, we noted a significant METTL3i-dependent IFN signalling in the ISG^High^ cell line OE21, despite its modest IR and RO induction. At the same time, it has been shown that innate immune responses to inverted Alus are enhanced by loss of the RNA editor ADAR1 (Ahmad et al., 2018), which acts on dsRNA and produces A to I edits that disrupt dsRNA structures (Cadena & Hur, 2019). We previously found that ADAR1 increased sensitivity to METTL3i in a CRISPR co-culture screen (Guirguis et al., 2023). The same screen identified the loss of genes associated with dsRNA sensing, antigen presentation and JAK–STAT signalling pathways to induce resistance to METTL3i, demonstrating the activating effect of the innate immunity system, in particular type I IFN signalling, on the cytotoxic T-cells, and generally on the adaptive immune system (Busselaar et al., 2024, Zitvogel et al., 2015).

RNA modifications, including m6A, play an important role in the ability of the innate immune system to distinguish ‘self’ from ‘non-self’ RNA, and thus prevent an inflammatory response against endogenous RNA species (Kariko et al., 2005, Durbin et al., 2016). m6A modification in particular may serve as a mark for endogenous mRNA to avoid recognition by RNA sensors. To this end, certain viruses were found to hijack the m6A writer complex to methylate their own RNA and promote immune evasion (Williams et al., 2019, Lu et al., 2021, Winkler et al., 2019). Others have suggested that loss of m6A activates the expression or enhances the stability of endogenous retroviral (ERV) elements (Chelmicki et al., 2021, Xu et al., 2021), which then elicit responses from RNA sensors. In our experiments, we confirmed the upregulation of transcripts encoding for ERV and other types of TE. However, these were not independent transcripts but sequences contained within aberrant transcripts that span long stretches of intronic and intergenic regions, where TEs are very common (Wells & Feschotte, 2020).

Although in some cancer cells METTL3 inhibition can be directly cytotoxic, this is not a prerequisite for its efficacy in more complex systems. For example, SKOV3 cell viability is only modestly affected by STC-15 when grown in monoculture. In contrast, when co-cultured with PBMC, SKOV3 cells are highly sensitive to STC-15 induced cell killing. These data suggest that cancer cell intrinsic transcriptomic changes that lead to type I IFN and ISG production can activate PBMC and enhance their cytotoxic activity. Interestingly, cancer and immune cells respond differently to the loss of m6A and RNA misprocessing, with type I IFN responses occurring mainly in the cancer cells (Fig. 8D). This is therefore a unique mode of action that differs from other IFN inducers such as STING or TLR agonists.

An intriguing finding from this study is that m6A modification levels differ across cell lines and correlate with both the magnitude of IR and RO events and the degree to which the type I IFN response is induced. This suggests that cancers with higher m6A modification levels are more likely to be responsive to STC-15. While METTL3i monotherapy can exert profound anti-tumour effects in some mouse models (Yankova et al. 2021), in other models co-dosing with ICI enhances efficacy and can result in complete regression of tumours. This strategy is currently being evaluated in a Ph1b/2 clinical trial (NCT06975293), where STC-15 is given in combination with toripalimab, a novel next-generation PD-1 inhibitor, in patients with solid cancer.

## Methods

### Cell lines

A549 (86012804), COV644 (07071908), OE21 (96062201) and SKOV3 (91091004) were purchased from ECACC. Caov3 (HTB-75) and EFO21 (ACC 235) were purchased from ATCC and DSMZ, respectively. All cell lines were incubated at 37°C and 5% CO_2_. Cell lines were verified to be negative for Mycoplasma contamination.

### GLORI m6A sequencing

GLORI (glyoxal and nitrite-mediated deamination of unmethylated adenosines) was carried out as described in Liu et al., 2023. Total RNA was purified by adding Trizol directly to cultured cells and then extracting RNA using Qiagen miRNeasy columns according to the manufacturer’s instructions including a DNase treatment step on column. Polyadenylated RNA was purified from 7.5 μg total RNA using the NEBNext PolyA mRNA magnetic isolation module (NEB E7490) according to the manufacturer’s instructions but eluting in 18 μl nuclease-free water and fragmented by incubating (8 min, 94 °C) with 2 μl 10x NEBNext RNA fragmentation buffer. The fragmentation reaction was stopped by adding 2 μl 10x NEBNext RNA fragmentation stop solution and RNAs were purified using an RNA clean and concentrator column (Zymo, R1015) eluting in 14 μl. 1 ng of spike-in probe mix consisting of 5 different sequences each with an m6A modification site modified to 5, 20, 50, 80 or 100 % was added to each sample. To convert unmodified A positions to Gs, RNA was first incubated (30 min, 50 °C) with 6 μl glyoxal solution (8.8 M in H2O, Sigma, 50649-25 ml) and 20 μl DMSO and then incubated (30 min, 50 °C) with 10 μl saturated H3BO3 and finally incubated (8h, 16 °C) with 50 μl deamination buffer (750 mM NaNO2, 40 mM MES pH 6, 10 μl glyoxal solution [8.8 M in H20]). RNAs were precipitated by incubating (overnight, -80 °C) with 300 μl ethanol and 1.5 μl Glycoblue (ThermoFisher, AM9515) and centrifuging (20000g, 30 min, 4°C). RNA pellets were washed, dried, incubated (10 min, 95°C) with 50 μl deprotection buffer (25 μl 1 M TEAA pH 8.6, 22.5 μl formamide) and purified using an RNA clean and concentrator column. RNA was incubated (30 min, 37 °C) with 5 μl T4 PNK buffer, 5 μl ATP, 1 μl Superase.In, 2 μl T4 PNK in a final 50 μl volume. RNAs were purified using an RNA clean and concentrator column, eluting in 6 μl, and generated into libraries for sequencing using the NEBNext multiplex small RNA library kit for Illumina according to the manufacturer’s instructions. Final libraries were pooled and size-selected by gel electrophoresis selecting for bands from 160-250 bp. Libraries were sequenced to a depth of greater than 100 million PE150 reads on a NovaSeq X plus Illumina machine. Data from two biological replicates were collected for each condition: A549, CAOV3, COV644, EFO21, OE21 and SKOV3 cells treated with DMSO vehicle or 1 μM STM3675.

### GLORI m6A analysis

Read 1 fastq files of paired end reads were trimmed of adapters using cutadapt (-O 10 -m 20 – trimmed-only -a AGATCGGAAGAGCACACGTCTGAACTCCAGTCAC) (Martin, 2011). Reads with greater than 3 As were removed and As in remaining reads (indicative of m6A modification) were converted to Gs and aligned using the STAR aligner (--outFilterMultimapNmax 10 --outFilterMismatchNmax 2) (Dobin et al., 2013) to a STAR index generated from the A-to-G converted sequences of the positive and reverse complemented negative strands of the GRCh38 genome and ensembl gene annotations. Reads were also aligned using the STAR aligner (--outFilterMultimapNmax 1 --outFilterMismatchNmax --alignIntronMin 1 --alignIntronMax 1 --alignEndsType EndToEnd) to a STAR index generated from A-to-G converted spike-in sequences. Previously converted A-to-G positions in aligned reads were converted back to As and the number of As and Gs were counted at each A position in the mRNA and lncRNA transcriptome using bcftools mpileup. The m6A proportion (A count/(A + G count)) at each site was then calculated. For each comparison the mean m6A proportion and mean A + G count across both replicates has to be greater or equal to 0.25 and 20, respectively, across all samples being compared. “MANE Select” canonical transcript exon annotations were used to determine the m6A levels in particular exon types.

### RNA-seq

RNA from *in vitro* cell line samples (A549, CAOV3, COV644, EFO21, OE21 and SKOV3) treated with DMSO vehicle or 1 μM STM3675 were generated into RNA-seq libraries by Lexogen using Ribocop rRNA-depletion and CORALL Total RNA-seq library generation and sequenced to a depth of greater than 100 million SE100 reads. RNA from an *in vitro* CAOV3 cell line treated with a range of doses of STC-15 and *in vivo* whole blood and tumour samples were generated into RNA-seq libraries by Novogene using their rRNA-depletion strand-specific sequencing service. Blood samples were also depleted for Globin RNA. Libraries were sequenced to a depth of greater than 40 million PE150 reads on a NovaSeq X plus Illumina machine. Data from three or five biological replicates were collected for each *in vitro* or *in vivo* condition, respectively.

### RNA-seq processing

Paired-end RNA-seq fastq files were trimmed of adapters using Trim Galore (Krueger et al., 2023) and then aligned to either the mouse GRCm39 or the human GRCh38 genome using the STAR aligner (Dobin et al., 2013). Unique molecular identifiers (UMIs) were extracted from single-end RNA-seq fastq files. Fastq files were then trimmed of adapters using cutadapt (Martin, 2011) according to Lexogen’s recommendations and aligned to the human GRCh38 genome using the STAR aligner (Dobin et al., 2013). Bam files were deduplicated using UMI tools (Smith et al., 2017) (--method=unique) and reads were converted back to randomised fastq files for Salmon gene counting (Patro et al., 2017). Normalised bigwig tracks were generated by dividing the signal at each nucleotide by the total genome signal and multiplying by 1 billion.

### Gene expression analysis

Salmon (Patro et al., 2017) was used to count the number of reads aligning to each gene. DESeq2 (Love et al., 2014) was then used to determine the shrunken log2 fold-change in RNA level and associated adjusted p-value (padj) for each METTL3 inhibitor-treated condition relative to its vehicle or DMSO-treated control. Pathway enrichment analysis was applied to significantly (padj < 0.05) up or down-regulated groups of genes using the clusterProfiler package (Xu et al., 2024) and gene ontology biological process gene sets.

### Transcriptional Run-On (RO) analysis

To define precisely RO regions, a gtf file consisting of 1000 nt chunks extending downstream of the 3’ end of each MANE Select-labelled GRCh38 human genome transcript up until the chunk before the 5’ end of the next transcript or 100000 nt, if there was no downstream gene in this region, was generated. For RO analysis in cell lines treated with STM3675, featureCounts (Liao et al., 2014) was then used to count reads aligning to each chunk. For each gene, run-on regions were determined from the first chunk to the chunk before two consecutive chunks below a read count of 5 either across any CAOV3 STM3675-treated samples or across any of the 6 cell line STM3675-treated samples, filtering out RO regions shorter than 10000 nt. Counts for each RO region were then determined using featureCounts and combined with Salmon gene counts.

DESeq2 (Love et al., 2014) was used to determine the log2 fold-change in RNA level and associated adjusted p-value for each METTL3 inhibitor-treated condition relative to its vehicle or DMSO-treated control. Significant RO events were defined as those with a log2 fold-change > 1 and padj < 0.05. To limit the possibility that increased gene expression may lead to some RO, genes with RO that also had a log2 fold-change > 1 were excluded. Significant RO regions defined for the CAOV3 STM3675-treated samples were used to analyse run-on in CAOV3 STC-

15-treated samples. For these samples, featureCounts was used to count read pairs aligning to these regions before determining the log2 fold-change in RO at these regions. For RO event analysis in *in vivo* samples, featureCounts was used to count read pairs aligning to regions spanning up to 20000 nt or until the 5’ end of the downstream transcript of “Ensembl canonical”-labelled protein-coding GRCm39 mouse genome transcripts. Significant RO events cell line samples were defined as those with a log2 fold-change > 1 and padj < 0.05, excluding those for which the gene with the RO had a log2 fold-change > 1. A more stringent padj < 0.00001 threshold was used to define significant RO events in *in vivo* samples.

### Intron Retention (IR) analysis

featureCounts (Liao et al., 2014) was used to count reads or read-pairs aligning to introns within MANE Select-labelled GRCh38 human genome transcript or “Ensembl canonical”-labelled protein-coding GRCm39 mouse genome transcripts. The log2 fold-change in RNA level and associated adjusted p-value for each intron relative to other introns in the gene, to control for increased intronic signal solely as a result of increased gene expression, using DEXseq (Anders et al., 2012) and relative to all other introns using DESeq for each METTL3 inhibitor-treated condition relative to its vehicle or DMSO-treated control was calculated. Significant IR events were defined as those with a DEXseq log2 fold-change > 0.5 and padj < 0.05 and DESeq log2 fold-change > 0 and padj < 0.05 and with intron width > 250 nt. A more stringent DEXseq padj < 0.00001 threshold was used to define significant RO events in *in vivo* samples. For all plots, IR events were further filtered for those downstream of long exons ≥ 300 nt. The DEXseq log2 fold-changes were used in all plots.

### J2 dsRNA immunoprecipitation (dsRIP)

J2 IP method was adapted from Gao et al., 2021. Caov3 cells were treated with either 1 μM STM3675 or DMSO for 24 h before collection, washing with PBS and pelleting.

For J2 RIP from whole cell RNA: RNA was purified with TRIzol reagent (Invitrogen 15596018) using the miRNeasy Mini Kit (QIAGEN) including DNase I treatment, according to manufacturer’s instructions and eluted in 60 μl RNase-free water. RNA was then split into input, J2 IP and IgG IP. For IP samples, RNA was diluted to 900 μl in dsRIP buffer (100 mM NaCl, 50 mM Tris-HCl (pH 7.4), 3 mM MgCl2, 0.5% IGEPAL-CA-630, cOmplete EDTA-free Protease inhibitor cocktail table (Roche), SUPERase-In RNase Inhibitor (Invitrogen AM2696)), then 5 μg of either anti-dsRNA J2 antibody (Jena Bioscience, RNT-SCI-10010) or mouse IgG control were added to the RNA. Tubes were incubated for 2 h with rotation at 4 °C. Protein G Dynabeads (Invitrogen 10003D) were washed in dsRIP buffer and then added to the RNA/antibody solution. Tubes were incubated for 1 h with rotation at 4 °C. Beads were then washed 4 times with ice cold 500 μl dsRIP buffer. RNA was then extracted directly from the beads with 1 ml TRIzol Reagent. 200 μl chloroform was added and mixed well before centrifuging for 15 min at 13000g at 4 °C. The upper aqueous phase was collected and a further 200 μl chloroform was added and the centrifugation was repeated. The upper aqueous phase was collected and an equal volume of isopropanol was added alongside 1 μl GlycoBlue Coprecipitant (Invitrogen AM9515). Tubes were inverted 10 times and then incubated at -20 °C for 20 min. Samples were centrifuges at 8000g for 10 min at 4 °C. Supernatant was discarded and pellet was washed with 800 μl of cold 75% ethanol. RNA was air-dried for 5 mins and then resuspended in 15 μl RNase-free water and concentration was measured with High Sensitivity RNA ScreenTape Assay (Agilent Technologies). Experiments were performed in duplicate.

For J2 RIP from cytoplasmic lysate: cells were lysed in 900 μl dsRIP buffer and centrifuged at 13000 rpm for 10 min to pellet debris. Supernatant was collected and divided for J2 IP and IgG IP. 5 μg of either J2 antibody or mouse IgG control were added to the cell lysates. Tubes were incubated for 2 h with rotation at 4 °C. Protein G dynabeads were washed in dsRIP buffer and then added to the RNA/antibody solution. Tubes were incubated for 1 h with rotation at 4 °C. Beads were then washed 4 times with ice cold 500 μl dsRIP buffer. RNA was then extracted directly from the beads with 700 μl Trizol reagent. 140 μl chloroform was added and mixed well before centrifugation for 15 min at 13000g at 4 °C. The upper aqueous phase was collected and 1.5 volumes 100% ethanol was added. The samples were applied to RNeasy Mini (QIAGEN) spin column and RNA purification for performed according to manufacturer’s instructions, including DNase treatment. RNA was eluted in 30 μl RNase free water and quantitated with High Sensitivity RNA ScreenTape Assay (Agilent Technologies). Experiments were performed in duplicate. IgG IPs did not return sufficient RNA to generate sequencing libraries.

### MDA5 RNA immunoprecipitation (MDA5 RIP)

Caov3 cells were treated with either 1 μM STM 3675 or DMSO for 24 h before collection, washing with PBS and pelleting. Cells were lysed in 900 μl dsRIP buffer and centrifuged at 13000 rpm for 10 min to pellet debris. Supernatant was collected and divided for MDA5 IP and IgG IP. 2.3 μg of either MDA5/IFIH1 antibody (Proteintech 21775) or rabbit IgG control were added to the cell lysates. Tubes were incubated for 2 h with rotation at 4 °C. Protein A Dynabeads (Invitrogen 10001D) were washed in dsRIP buffer and then added to the RNA/antibody solution. Tubes were incubated for 1 h with rotation at 4 °C. Beads were then washed once with ice cold 500 μl dsRIP buffer. Beads were resuspended in 500 μl dsRIP buffer and 1 μl RNase T1 (Thermo Scientific EN0541) and incubating for 30 min with rotation at 4 °C. Beads were then washed 3x with 500 μl dsRIP buffer.

RNA was then extracted directly from the beads with 700 μl Trizol reagent. 140 μl chloroform was added and mixed well before centrifugation for 15 min at 13000g at 4 °C. The upper aqueous phase was collected and 1.5 volumes 100% ethanol was added. The samples were applied to RNeasy Mini (QIAGEN) spin column and RNA purification for performed according to manufacturer’s instructions, including DNase treatment. RNA was eluted in 30 μl RNase free water and quantitated with High Sensitivity RNA ScreenTape Assay (Agilent Technologies). Experiments were performed in duplicate. IgG IPs did not return sufficient RNA to generate sequencing libraries.

### dsRIP-seq and RIP-seq library preparation

Libraries for input and IP samples were generated using the NEBNext UltraExpress RNA Library Prep Kit (NEB), largely according to manufacturer’s instructions for rRNA-depleted samples, with the following specified exceptions. First, rRNA was depleted from samples using rRRR, according to the published protocol (Singh et al., 2024). After DNase I digestion, RNA was purified using Agencourt RNAClean XP beads and eluted in Fragmentation Master Mix according to the NEBNext UltraExpress RNA Library Prep Kit protocol. RNA was fragmented at 94 °C for 12 min, except for the MDA5 IP libraries for which a 4 min fragmentation was used. Finally, libraries were quantitated on a D100 DNA ScreenTape Assay (Agilent Technologies) and sequenced by Illumina, PE150.

### IP-seq data analysis

Reads were trimmed with cutadapt (Martin, 2011). For MDA5 IP (and corresponding cytoplasmic input RNAseq), rRNA reads were removed using bbduk version = 39.01 (Bushnell., 2018). Reads were aligned with STAR version = 2.7.10b (Dobin et al., 2013) and to the human reference genome GRCh38. Read count tables were generated with featurecounts (Liao et al., 2014) using the Ensembl primary annotation release 108, TE annotations generated from UCSC RepeatMasker (Jin et al., 2015), or our own annotation of run-on and intron retention coordinates described above. Read counts from our defined run-on and intron retention events were appended to the primary annotation count table. Differential expression analysis was performed with edgeR (Chen et al., 2024) using single and double contrasts (IP vs input for each treatment, IP vs IP, input vs input, (IP vs input) vs (IP vs input)). Inverted Alus were considered to be any pair of Alus on opposite strands with a maximum separating distance of 1 kb.

### Co-culture Tumour-Killing Assay

The co-culture studies have been performed at Charles River Laboratories. SKOV3-NucLight Red cells were co-cultured with donor PBMC, with STC-15 or DMSO control treatments added at the point of co-culture at the indicated concentrations. Co-culture plates were incubated at an IncuCyte S3 System, and images were collected every 2 h for 72 h. SKOV-NLR cell growth was calculated based on the red fluorescence channel, quantified as red object count per image, and normalised as a percentage to *T* = 0 h. SKOV-NLR apoptosis was determined by detecting caspase-3/7 signal in the green fluorescence channel colocalized with SKOV-NLR signal detected in the red fluorescence channel. Each treatment condition was repeated in triplicate wells. Total of 3 donors were tested. At 72h, the triplicate well culture supernatants were combined and frozen for cytokine analysis (see below).

For 3D spheroids generation, SKOV3-NLR cells were seeded in ultra-low attachment plates, centrifuged, treated with IFNg as above and left for 48 h to form the spheroid. At 48h, PBMC and STC-15 or DMSO control treatments, prepared as above, were added. Co-cultures were placed in the IncuCyte and imaged every 2 h for 7 days. Total of 3 donors were tested. Each experimental condition was tested in quadruplicate wells. Image analysis was performed using IncuCyte’s Spheroid Module (Sartorius). Briefly, SKOV-NLR signal was detected in the red fluorescence channel and quantified within the boundary of the spheroid in the brightfield image. Data was normalised to 24h after initiation of co-culture, to allow for settling time after PBMC addition. PBMC migration at End point was quantified by creating a mask over the brightfield image with size exclusion filter to exclude the large central spheroid body.

### In vivo studies

MC38 studies were conducted at Sygnature Discovery, UK. All experiments were performed in accordance with the Animal Science Procedures Act (ASPA) 1986 (UK) and Establishment and Project licence guidelines.

MC38 cells were purchased from the NCI. Anti-PD1 (clone RMP1-14 antibody, lot number 810421N1) was purchased from Bio-X-Cell. STC-15 was dissolved in HPBCD/Sodium acetate buffer pH4.6, 50mM (10%:90%; w/v) at 10 mg/ml.

Female C57Bl/6 mice (Envigo UK), were implanted subcutaneously (s.c.) on the left flank with 1x10^7^ MC38 cells in serum free DMEM. Tumours were measured three times weekly by digital calipers.

#### MC38 efficacy study

Seven days post inoculation, animals were randomised onto study in groups (10 mice/group). Dosing groups were: (1) Vehicle (PO, BID) + PBS (IP, BIW); (2) STC-15 (15mg/kg PO, BID) + PBS (IP, BIW); (3) STC-15 (50mg/kg PO, BID) + PBS (IP, BIW); (4) Vehicle (PO, BID) + anti-PD1 (10mg/kg IP, BIW) (5) STC-15 (15mg/kg PO, BID) + anti-PD1 (10mg/kg IP, BIW) (6) STC-15 (50mg/kg PO, BID) + anti-PD1 (10mg/kg IP, BIW). Dosing started at the day following randomisation (Day 1) and continued for 14 days. Animals were monitored daily for clinical signs. Key observations included effects on mobility, food and water intake, body weight changes and tumour condition. At termination, tumours were cut into two pieces, for IHC and for gene expression analysis using Nanostring. For IHC (n=6 tumours/group), the tumour was put into 10% buffered formalin for 24 h then transferred to 70% ethanol. For Nanostring analysis (n=3 tumours/group), tumours were snap frozen in liquid nitrogen and stored at -80°C.

#### MC38 RNA-seq study

Mice were randomised into 7 groups (5 mice/group). Six groups were orally dosed with 100 mg/kg STC-15 once daily, for sample collection at T=0, T=1h, T=4h, T=12h, T= 48h and T=120h. One group was orally dosed once daily with vehicle control, for sample collection at T=120h. The mice in the T=0 group received 6 treatment doses and were terminated 24h after the 6^th^ dose. The mice in the other groups received 7 treatment doses and were collected at the indicated time following the 7^th^ dose. For RNA-seq, blood and tumour samples for were collected at termination. Up to 500 μL of blood was collected by cardiac puncture, stored in RNAprotect tubes (Qiagen, 76554) and left at RT. Tumours were cut in half, placed in cryovials and snap frozen.

**For additional Methods, Reagents used in this work, Supplementary Figures and Supplementary Files description, please refer to the Supplementary Information and Figures file.**

## Acknowledgement

The authors would like to thank all Storm Therapeutics’ colleagues, past and present, who contributed to the development of STC-15 directly and indirectly. The authors would like to acknowledge the invaluable contribution of our CRO collaborators, including but not limited to Evotec, Charles River Laboratories, Sygnature Discovery, Crown BioScience and Histologix. We thank Profs. Tom Cech (University of Colorado Boulder), Mark Dawson (Peter MacCallum Cancer Centre), Tony Kouzarides (University of Cambridge) and Stephanie Halene (Yale University) for critical review of the manuscript.

