## Supplementary Information and Figures for "Inhibition of METTL3 by STC-15 induces RNA misprocessing that results in dsRNA formation and activates innate immunity"

#### Additional Methods

##### Immunofluorescence staining

Caov3 cells were fixed with 4% paraformaldehyde for 10 min at RT, permeabilized with 0.2% Triton X-100 in PBS for 10 min and blocked with 10% donkey normal serum (AbCam, ab7475) in PBS for 2-3h RT. Cells were stained with the dsRNA-specific antibody J2 and with anti-IFIT1 antibody at 4°C overnight. Cells were incubated with secondary antibodies 2h RT in the dark (donkey anti-mouse-Biotin and donkey anti-rabbit-AF488) and then with Streptavidin-AF555 (Invitrogen,S32355, 1:400) for 2h at RT in the dark. Cells were stained with DAPI (1:5,000) for 10min at RT in the dark, before mounting with Prolong Gold Diamond Antifade Reagent (Invitrogen, P36970). Cells were gently washed 3 times between each incubation step. See Supplementary Table S1 for antibodies' details.

##### qPCR

To generate cDNA, total RNA from Caov3 cells was reverse-transcribed with the SuperScript VILO cDNA synthesis kit (Thermo Fisher) according to the manufacturer's instructions. Quantitative real-time PCR was performed on the Applied Biosystems QuantStudio 5 using TaqMan Fast Advanced Master Mix and TaqMan gene expression probes (Thermo Fisher), according to manufacturer's protocol. mRNA levels were normalised to the housekeeping gene PPIA. All samples were assayed in triplicate with relative quantification of target gene expression performed using the comparative cycle threshold (CT) method. TaqMan probes are listed in Supplementary Tables S2.

##### Westen Blot

Caov3 cells were harvested by trypsinization and lysed using RIPA buffer supplemented by protease and phosphatase inhibitors. Total protein (25 µg) was loaded on a 4%–12% Bis-Tris gel. For Western Blot to validate MDA5 IP, input, bound and unbound samples were loaded at volumes equivalent to 20 µg of input. Resolved proteins were then transferred to a nitrocellulose membrane and blocked for 1 h with 5% milk/TBST. Antibodies, as detailed in Supplementary Table S1, were diluted in 2.5% milk/TBST and incubated with the blocked membranes overnight at 4°C. Following 3 washes with TBST, the secondary antibody was incubated with the membrane for 1 h at RT. After 3 additional washes with TBST, signal was developed using electroluminescence.

##### Secreted cytokine assays

Caov3 cells were treated with STM3675 for 48 h. Tumor-conditioned media were collected, spun down at 800 × g for 2 min to remove cell debris, and stored at −20°C. IFNβ levels were assessed in duplicates using the S-PLEX Human IFNβ Kit (Meso Scale, K151ADRS) following the manufacturer's instructions. CXCL10 levels were assessed in duplicates using Quantikine ELISA, Human CXCL10 / IP-10 Immunoassay (R&D Systems, DIP100) according to the manufacturer's protocol.

Frozen supernatant samples from co-culture experiments were thawed at RT prior to analysis by Luminex Multiplex. Bio-Plex Pro Human Cytokine IL1b (Bio-Rad, #171B5001M), IFNg (Bio-Rad, #171B5019M), and TNFa (Bio-Rad, #171B5026M) analyte bead sets were prepared in Bio-Plex assay buffer (Bio-Plex Pro Reagent Kit III, #171304090M) and washed 2 times using wash buffer (Bio-Plex Pro Reagent Kit III, #171304090M) on the Bio-Plex Pro plate washer. Cell supernatant samples were incubated with analyte beads for 30 min at RT on a shaker at 850 rpm. Following incubation, 3 washes were performed and a detection antibody was applied. Samples were incubated on the shaker at 850 rpm for 30 min at RT. A further 3 washes were performed, and streptavidin-PE (Bio-Plex Pro Reagent Kit III, #171304090M) was added for 10 min at RT. A final 3 washes were performed, and beads were resuspended in assay buffer. Data were acquired on the Bio-Plex 200 System, which had been calibrated with a Bio-Rad Calibration Kit (Bio-Rad, #171203060).

***In vitro* T-cell proliferation**

Healthy Donor Human PBMC were isolated from two donors using density gradient centrifugation and stained with Cell Proliferation Dye (Invitrogen eBioscience, #15510597) according to the manufacturer's instructions. Cells were seeded in a 96 well round-bottom plate in assay media of RPMI + 10% Human AB serum, 20 mM HEPES, 4mM L-glutamine + 100 U/mL penicillin / streptomycin. The cells were pre-incubated with STC-15 at the indicated concentrations or with DMSO control for 24 h, after which anti-CD3 (5 µg/ml, Biolegend, 300332) + anti-CD28 (1 µg/ml, Biolegend, 302934) were added. Following 48 h stimulation, cells were stained for CD4 (Biolegend, 344604) and CD8 (Biolegend, 301028) and analysed by flow cytometry to assess viability and proliferation of CD4+ and CD8+ populations.

***In vivo* studies**

A20 studies were conducted at Crown Bioscience, UK. All studies were compliant with the UK Animals Scientific Procedures Act 1986 (ASPA) in line with Directive 2010/63/EU of the European Parliament and the Council of 22 September 2010 on the protection of animals used for scientific purposes.

A20 cells were purchased from ATCC (Cat. TIB-208).

CD8 depletion study. Female BALB/cN (Charles River) were implanted s.c. with 5 x $10^5$ viable cells in 0.1 ml PBS, injected into the left flank. Mice (n=10) were randomly allocated to the treatment groups. Treatment started the following day and was continued for up to 25 days. STC-15 was given PO at 100 mg/kg once daily. Anti-CD8 antibody (clone 2.43, 2B Scientific BioXcell, BP0061) was given by i.p. injection BIW while dosing continued. Tumour volumes were measured three times a week for the duration of the efficacy study. Body weights were monitored daily throughout the dosing phase.

STC-15 efficacy and re-challenge study. Female BALB/cN (Charles River) were implanted s.c. with 5 x $10^5$ viable cells in 0.1 ml PBS, injected into the left flank. Mice were randomly allocated to the treatment groups. Treatment started the following day and was continued for up to 19 days. STC-15 was given PO at 50 mg/kg BID. Anti-PD1 antibody (Crown Bioscience, Lot #1120L365) was given by i.p. injection BIW while dosing continued. Tumour volumes were measured three times a week for the duration of the efficacy study. Body weight was monitored daily throughout the dosing phase. The re-challenge phase of the study commenced at Study Day 61, following a four-week treatment-free observation phase during which tumour volume and body weight were monitored three times a week. Treatment naïve mice (n=6), or mice previously treated with STC-15 + anti-PD1 combination and experienced complete tumour regression (n=6) were re-inoculated on the right flank with fresh 5 x $10^5$ A20 cells in 0.1 ml PBS. There was no further treatment during the re-challenge phase. Tumours continued to be monitored three times weekly and body weight was monitored daily.

**Nanostring analysis on tumours**

Snap frozen mouse tissues were transferred to a Lysing Matrix D 2 ml tubes (MP Biomedicals 6913-500) containing 0.9 ml of QIAzol reagent (QIAGEN 79306) and disrupted by homogenization using Bead Mill 24 Homogenizer (Fisher Scientific). Total RNA was then extracted using the RNeasy Plus Universal Mini kit (QIAGEN 73404) on a QiaCube device (QIAGEN). RNA was quantified using Qubit RNA XR Assay Kit (Invitrogen Q33223) on a Qubit 4 Fluorometer (Invitrogen Q33226). RNA quality was assessed using RNA Screen Tapes (Agilent 5067-5576) on a 2200 Tape Station device (Agilent).

For gene expression analysis on the NanoString nCounter system, 100 ng of RNA was hybridized to a multiplexed nucleotide probe pool (Mouse Pan Cancer IO360 Panel) for 16 h at 65 °C. Enriched targets were purified and quantified using the nCounter MAX Analysis System. RCC files were analysed using nSolver Analysis Software (Version 4.0). Negative and positive controls were included in the Pan-Cancer IO-360 Panel probe sets. Samples were analysed using nSolver's advanced analysis module according to Nanostring's nCounter Advanced Analysis 2.0 user manual (MAN-10030-03).

**CD8+ infiltration by immunohistochemistry (IHC)**

Samples were processed to FFPE blocks, from which 4 µm sections were cut for H&E and CD8+ staining (using an anti-CD8, clone 4SM15 (E-Bioscience, 14-0808-82) antibody). The stained slides were digitally scanned, and cell based quantitative image analysis performed using the Indica Labs HALO® 3.0 image analysis platform, with subsequent spatial analysis of the infiltration of the positive cells across the tumour-stroma interface.

**Reagents Tables**

Table 1. Antibodies

| Antibody | Company | Dilution (WB/) | Dilution (IF) | Cat Number | Clone |
|---|---|---|---|---|---|
| p-STAT1 | CST | 1:1000 | | 9167S | 58D6 |
| STAT1 | CST | 1:1000 | | 9172 | |
| IFIT1 | CST | 1:1000 | 1:250 | 14769S | D2X9Z |
| IRF3 | CST | 1:1000 | | 4302S | D83B9 |
| p-IRF3 | CST | 1:1000 | | 37829 | E7J8G |
| ISG15 | CST | 1:1000 | | 2758S | 22D2 |
| OAS1 | CST | 1:1000 | | 14498S | |
| PKR | AbCam | 1:2000 | | ab184257 | EPR19374 |
| P-PKR | AbCam | 1:1000 | | ab32036 | E120 |
| IRF7 | CST | 1:1000 | | 4920 | |
| GAPDH | Sigma-Aldrich | 1:10000 | | G9545-100UL | |
| Donkey anti-rabbit (HRP) | Sigma-Aldrich | 1:10000 | | GENA934-1ML | |
| J2 (dsRNA) | SCICONS (2bscientific) | | 1:200 | 10010200 | |
| Donkey anti-mouse (Biotinylated) | AbCam | | 1:100 | ab7060 | |
| Donkey anti-rabbit (AF488) | Invitrogen | | 1:2,000 | A21206 | |
| dsRNA J2 (for IP) | Jena Bioscience | | | RNT-SCI-10010 | |
| MDA5/IFIH1 | Proteintech | | | 21775-1-AP | |
| Normal mouse IgG | Sigma-Aldrich | | | 12-371 | |
| Normal Rabbit IgG | CST | | | 2729S | |

Table 2. TaqMan™ probes

| Gene | Probe ID | Probe / Quencher |
|---|---|---|
| OAS1 | Hs00973637_m1 | FAM-MGB |
| ISG15 | Hs00192713_m1 | FAM-MGB |
| IFIT1 | Hs03027069_s1 | FAM-MGB |
| IFN-β | Hs01077958_s1 | FAM-MGB |
| PPIA | 4326316E | VIC-MGB |

Table 3. Other reagents

| Reagent | Company | Cat Number |
|---|---|---|
| Ruxolitinib | Enzo Life Science | ENZ-CHM156 |
| Bay-985 | Selleck Chemicals | S8935 |

Table 4. GLORI spike-in probe sequences

| | |
|---|---|
| spike in-5% | AGCGCAGCGAACAUGACACGUGCUCAACAGUGUAGUUGG(m6A)CUUCCUCGCAUCUAGCCGCUAUGCACGUGCAGCU |
| spike in-20% | AUCUACCUGUCCAGUAGCCUUCAGGAUCAUGCUGUCUGACUUGCUGG(m6A)CAUCAUUCUAGUGCCAUAACUUCAGC |
| spike in-50% | UAUGGUUUAGAUUGAAUGUUGUUGUUGUGAGG(m6A)CCUUAUGUAAGUUGAAUGGUAAUAGGAAGAAGUUUG |
| spike in-80% | UUGUUUGAUGGAGGGGGAUAAUUAUUGGA(m6A)CAGGUAGUUAAUAUUGUAUAAUGUUGUAAGAUUAAAGAGGG |
| spike in-100% | GGAUUAGUUAGUAGGUGGGGUAAUGGUUUAUGA(m6A)CCUGAUGAUUUUUAGUUGGUUUGAGAGGAUGAUUAGU |

#### Supplementary Files

**Supplementary Table 1**
Significant IR and RO events identified in Caov3 cells.

**Supplementary Table 2**
Over-represented gene ontology biological process (GO:BP) terms for genes with significant IR events or RO events in Caov-3 cells.

**Supplementary Table 3**
Significant IR and RO events identified in any cell line (A549, CAOV3, COV644, EFO21, OE21, SKOV3). STM3675 treated samples compared to DMSO control treated samples.

**Supplementary Table 4**
Enrichment analysis of dsRIP-seq (J2 antibody). dsRIP compared to input RNA-seq, dsRIP (STM3675) compared to dsRIP (DMSO), input RNA-seq (STM3675) compared to input RNA-seq (DMSO), and "double-strandedness" double contrast.

**Supplementary Table 5**
Enrichment analysis of dsRIP-seq (J2 antibody) from cytoplasmic input samples. dsRIP compared to cytoplasmic input RNA-seq, dsRIP (STM3675) compared to dsRIP (DMSO), cytoplasmic input RNA-seq (STM3675) compared to cytoplasmic input RNA-seq (DMSO).

**Supplementary Table 6**
Transposable elements in IR and RO events in Caov3 cells.

**Supplementary Table 7**
Enrichment analysis of MDA5 RIP-seq from cytoplasmic input samples. MDA5 RIP compared to cytoplasmic input RNA-seq, MDA5 RIP (STM3675) compared to MDA5 RIP (DMSO)

**Supplementary Table 8**
Differential gene expression in cell lines (A549, CAOV3, COV644, EFO21, OE21, SKOV3), Caov3 dose response and MC38 mouse model (blood and tumour)

**Supplementary Table 9**
Significant IR and RO events identified at any time point in either blood or MC38 tumour. STC-15 treated samples at time points after the last dose compared to vehicle control treated samples

**Supplementary Video 1**
SKOV3 spheroids co-cultured with donor PBMC, treated with DMSO control.

**Supplementary Video 2**
SKOV3 spheroids co-cultured with donor PBMC, treated with 1 μM STC-15.

### Supplementary Figure 1.

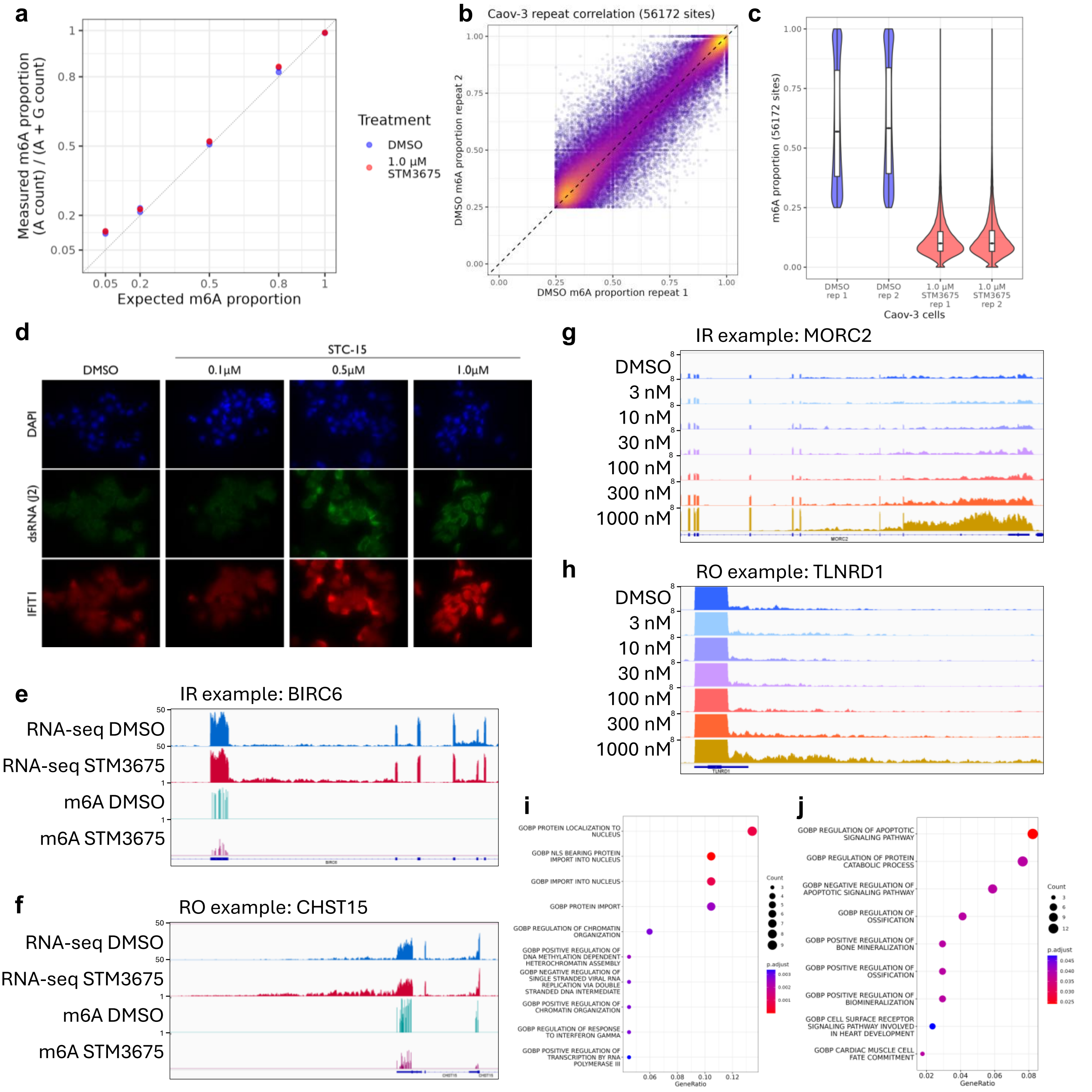


**Supplementary Figure 1. (a)** Scatter plot showing the measured vs expected m6A proportion at m6A sites in spike-in probes within each GLORI sample **(b)** Scatter plot comparing the proportion of m6A modification at sites in Caov3 cells treated with DMSO between two replicates. Points are coloured from high (yellow) to low (purple) density. **(c)** Violin and box and whisker plots showing the distribution of m6A proportions in Caov3 cells treated with DMSO or 1 µM STM3675 (2 replicates for each condition) **(d)** Immunofluorescence images showing dsRNA and IFIT1 protein levels, as detected by the J2 and anti-IFIT1 antibodies, respectively, in Caov3 cells treated with DMSO or 0.1, 0.5 or 1.0 µM STC-15. **(e-f)** IGV tracks showing RNA-seq and m6A proportions (GLORI data) from Caov3 cells treated for 24 h with either DMSO or 1 µM STM3675, exemplifying an intron retention (IR) event **(e)** and a run-on (RO) event **(f)**. **(g-h)** IGV tracks showing RNA-seq from Caov3 cells treated for 24 h with either with DMSO or 3, 10, 30, 100, 300 and 1000 nM STC-15, exemplifying an IR event **(g)** and an RO event **(h)**. **(i-j)** Dot plots showing enriched gene ontology biological process (GO:BP) terms for genes showing significant IR **(i)** or RO **(j)** events in Caov3 cells upon treatment with 1 µM STM3675. Each dot represents a GO term, with its position on the x-axis indicating the gene ratio (number of input genes associated with the term divided by the total number of input genes). Dot size reflects the count of genes mapped to each term, while color intensity corresponds to the adjusted p-value, indicating statistical significance of enrichment.

### Supplementary Figure 2.

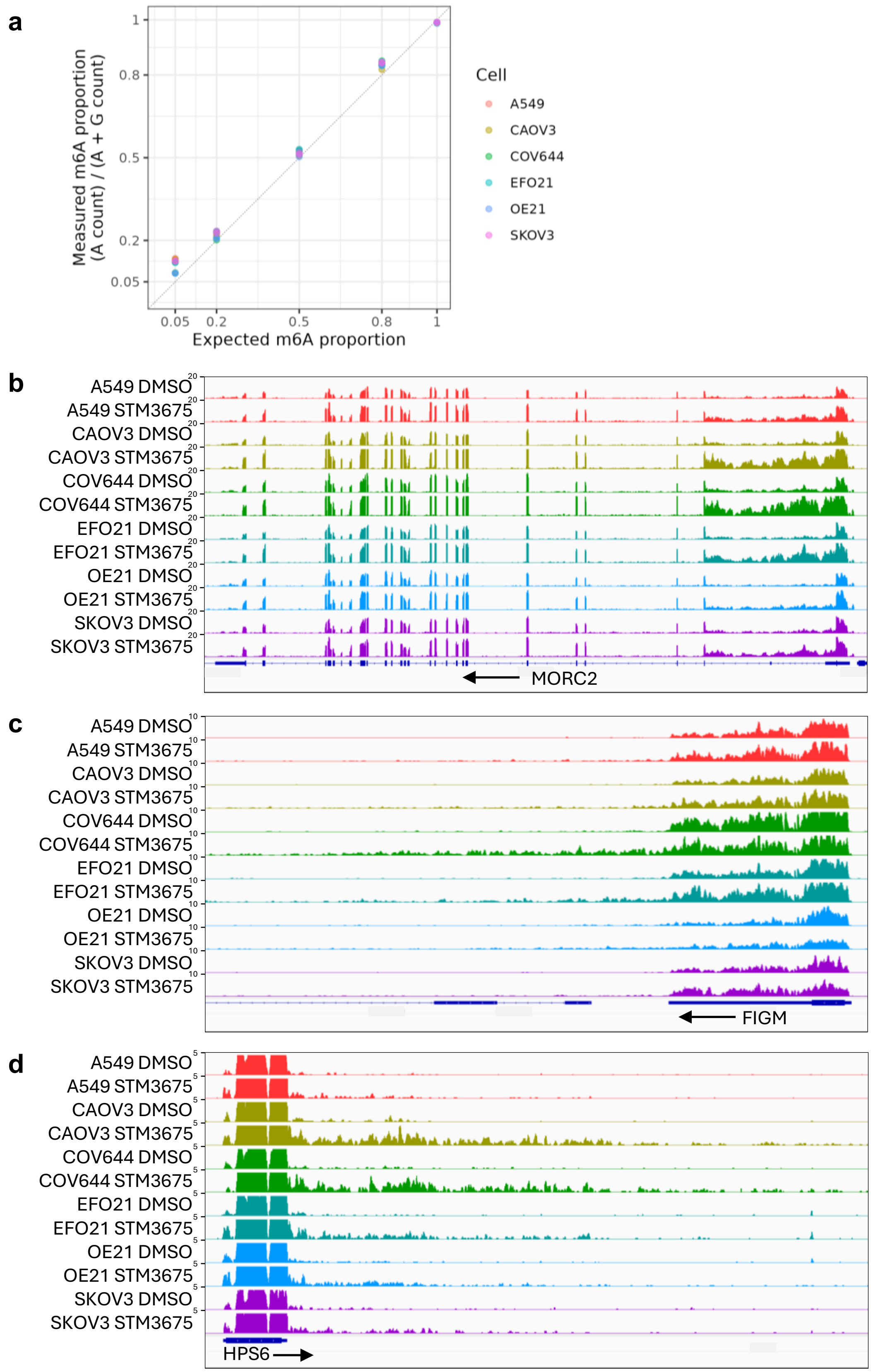


**Supplementary Figure 2.** Examples of IR/RO events across 6 cell lines. **(a)** Scatter plot showing the measured vs expected m6A proportion at m6A sites in spike-in probes within each GLORI sample **(b-d)** IGV tracks showing RNA-seq from 6 cell lines treated for 24 h with either with DMSO or 1 μM STM3675 exemplifying an IR event **(b)** and two run-on (RO) events **(c-d)**.

### Supplementary Figure 3.

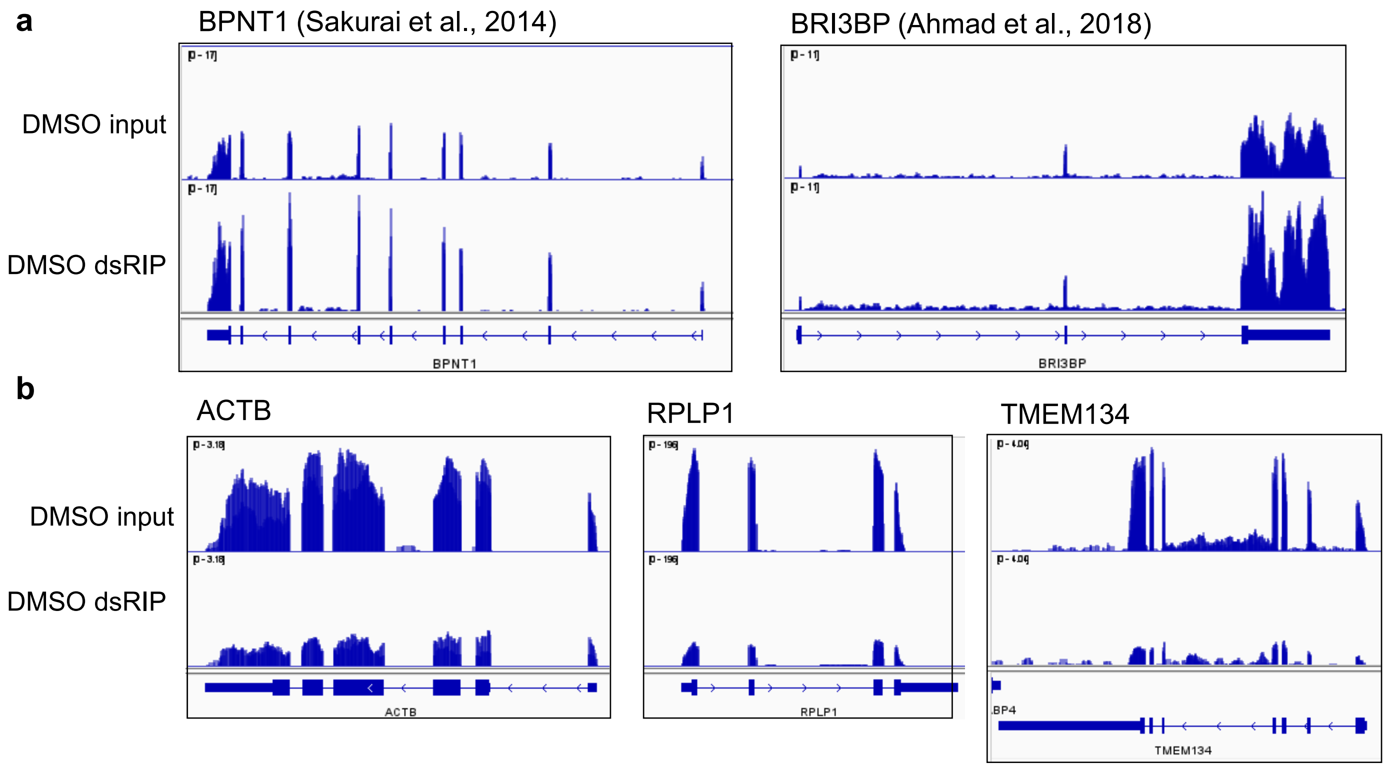


**Supplementary Figure 3.** Validation of dsRIP-seq – positive and negative controls. IGV tracks for input (RNA-seq) and dsRIP-seq from Caov3 treated for 24 h with DMSO (two replicates overlayed) are presented. Tracks are normalised to library size. **(a)** Positive controls selected from the literature to be expected to form double stranded RNA are enriched by dsRIP. **(b)** House-keeping genes are not enriched by dsRIP, despite high expression levels.

### Supplementary Figure 4.

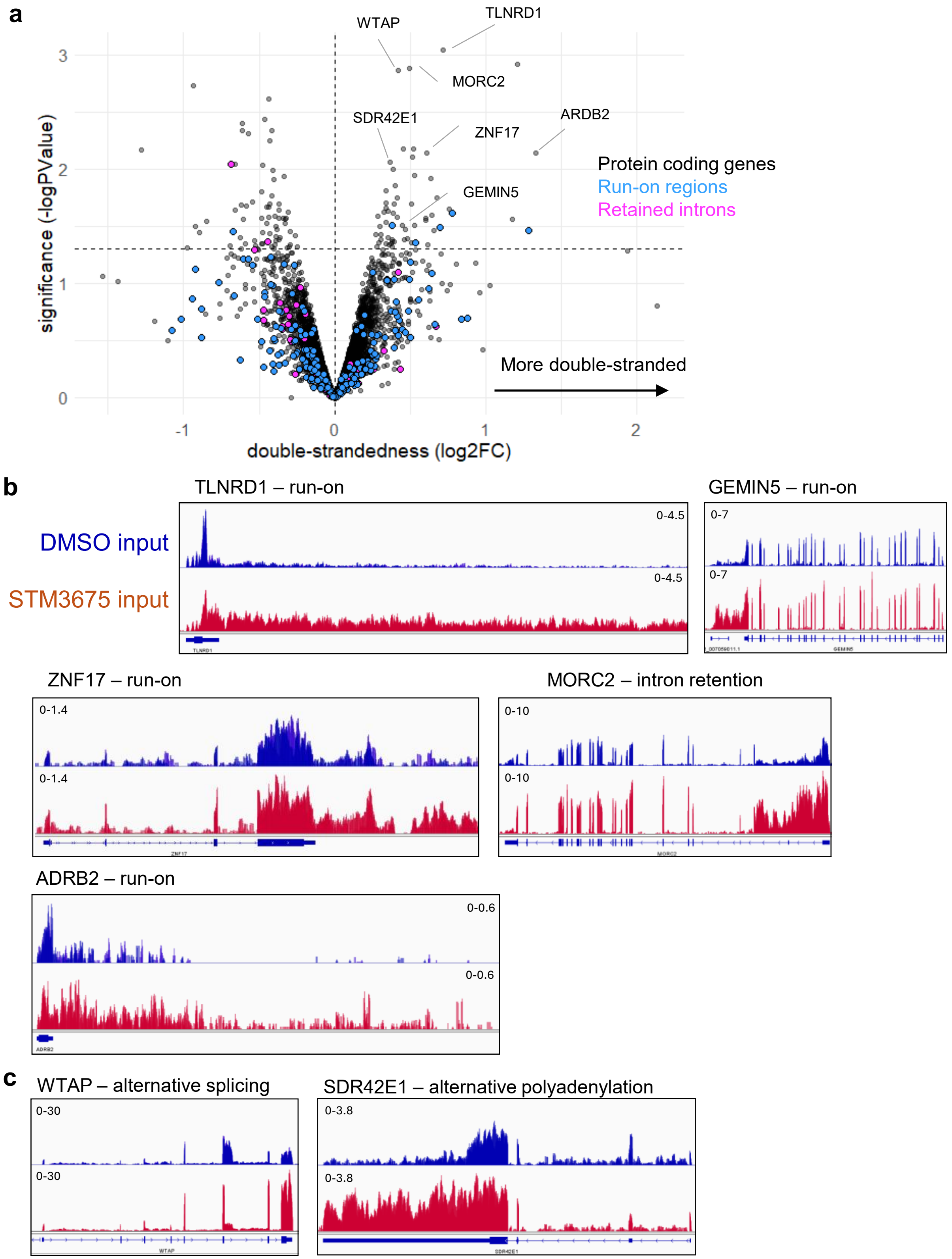


**Supplementary Figure 4.** Few transcripts show a change in 'double-strandedness' upon STM3675 treatment. **(a)** A double contrast in edgeR is used to compare enrichment of each species (dsRIP vs input) upon STM3675 treatment (STM3675 vs DMSO). Only RNA species that show a significant (FDR < 0.05) change in dsRIP in STM3675 vs DMSO are displayed. Canonical transcripts are shown in grey, IRs in pink, and ROs in blue. **(b-c)** Normalised input (RNA-seq) IGV tracks for DMSO and STM3675-treated Caov3 cells. Many of the RNA species showing a change in double-strandedness are the canonical transcripts of genes which experience either an IR or RO event **(b)** or an alternative splicing or polyadenylation event **(c)**.

### Supplementary Figure 5.

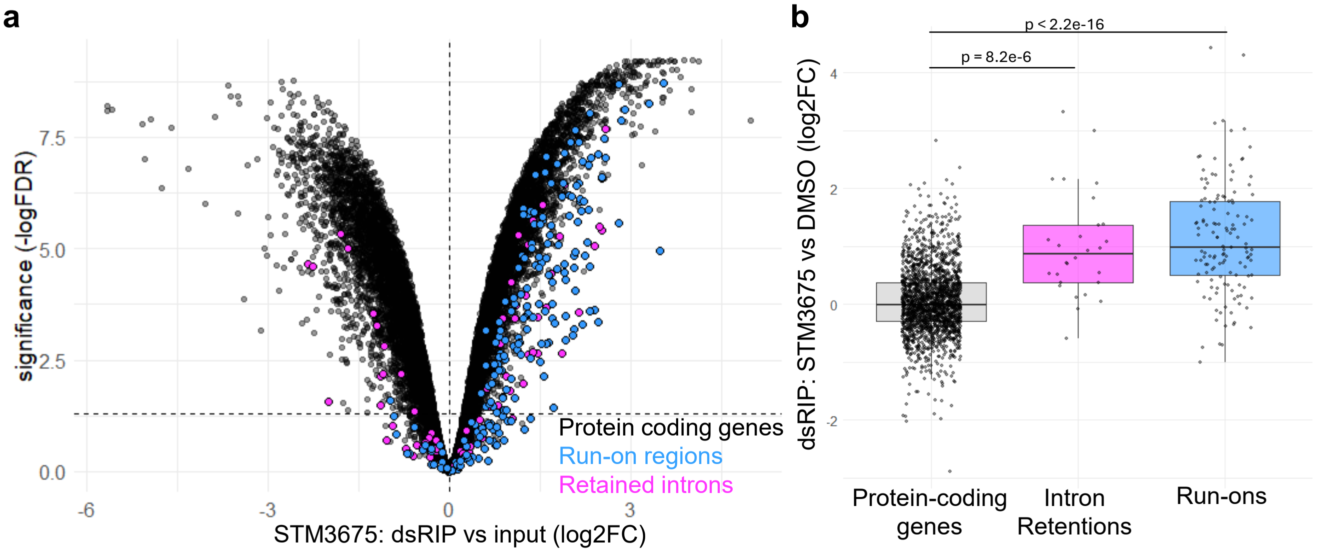


**Supplementary Figure 5.** Cytoplasmic dsRIP-seq. dsRIP experiment on the cytoplasmic fraction of Caov3 treated for 24 h with 1 µM STM3675 or DMSO. **(a)** Enrichment of RNA species in the J2 dsRIP-seq compared to input. Canonical transcripts are shown in grey, IRs in pink, and ROs in blue. **(b)** Box and whisker plot of the fold-change in dsRIP pulldown after 24 h treatment with 1 µM STM3675 vs DMSO. Only species that are classed as dsRNA in either DMSO or STM3675 are included (dsRIP vs input: log2FC > 1, FDR < 0.05). Canonical protein coding transcripts, IRs and ROs are displayed separately.

### **Supplementary Figure 6.** MDA5 RIPseq validation

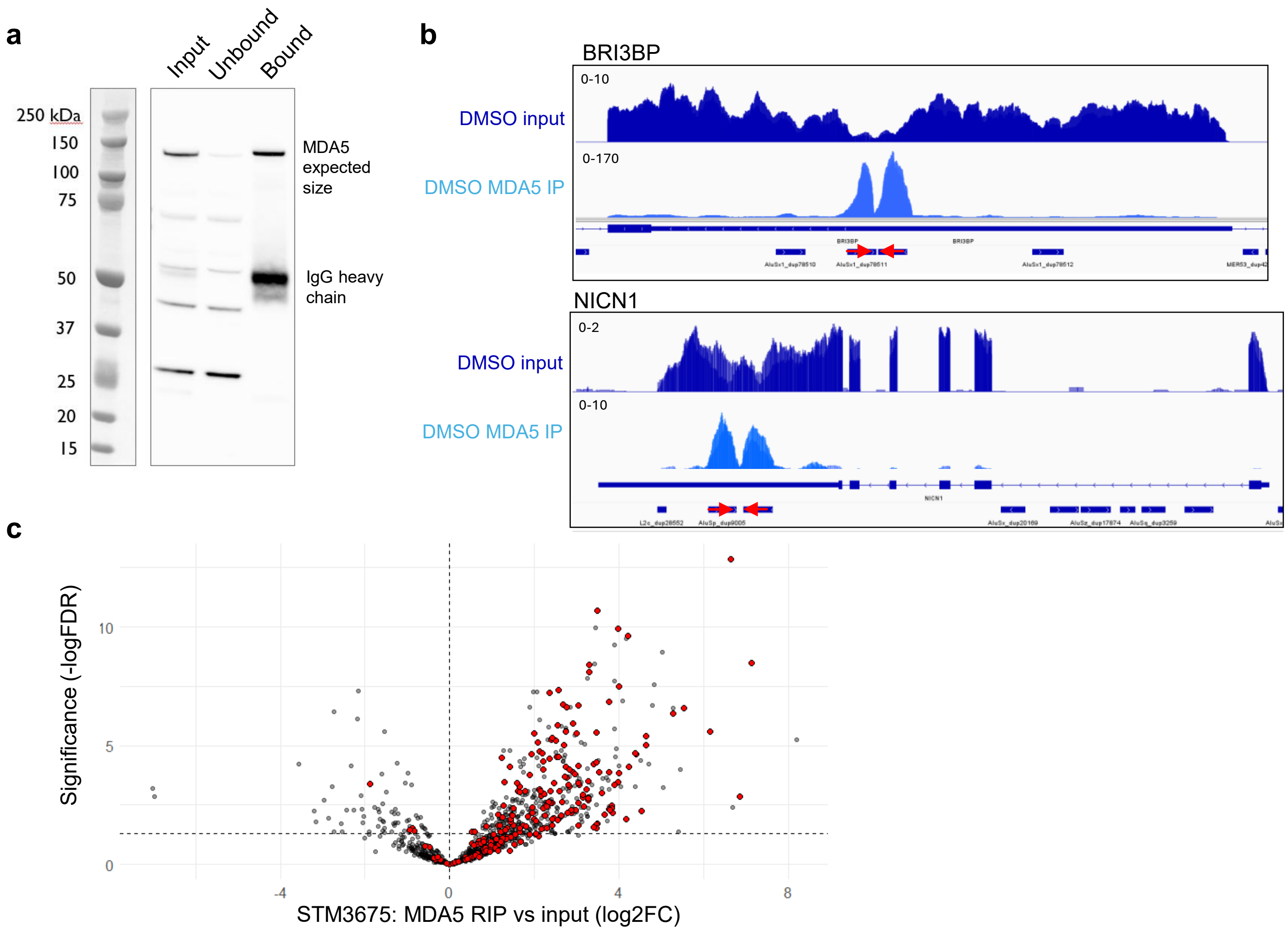


**Supplementary Figure 6.** MDA5 RIPseq validation. **(a)** anti-MDA5 Western blot of cytoplasmic input, unbound and bound fractions of anti-MDA5 IP from Caov3 cells treated with DMSO for 24 h. There is enrichment of MDA5 in the bound fraction, and depletion from the unbound fraction after anti-MDA5 IP. Non-specific bands of the wrong size detected by the anti-MDA5 antibody by Western blot are not enriched by the immunoprecipitation, while the non-crosslinked antibody is released from the beads in the bound fraction. **(b)** IGV tracks for showing MDA5 IP and cytoplasmic input of two genes, BRI3BP and NICN1, reported to form dsRNA in their 3' UTRs. Red arrows show inverted Alu pairs, note the differences in scaling for each track. **(c)** MDA5 IP vs input in STM3675 treated Caov3 cells, grey points are all transposable elements within IR/ROs, red points identify Alus that are part of an inverted Alu pair.

### Supplementary Figure 7.

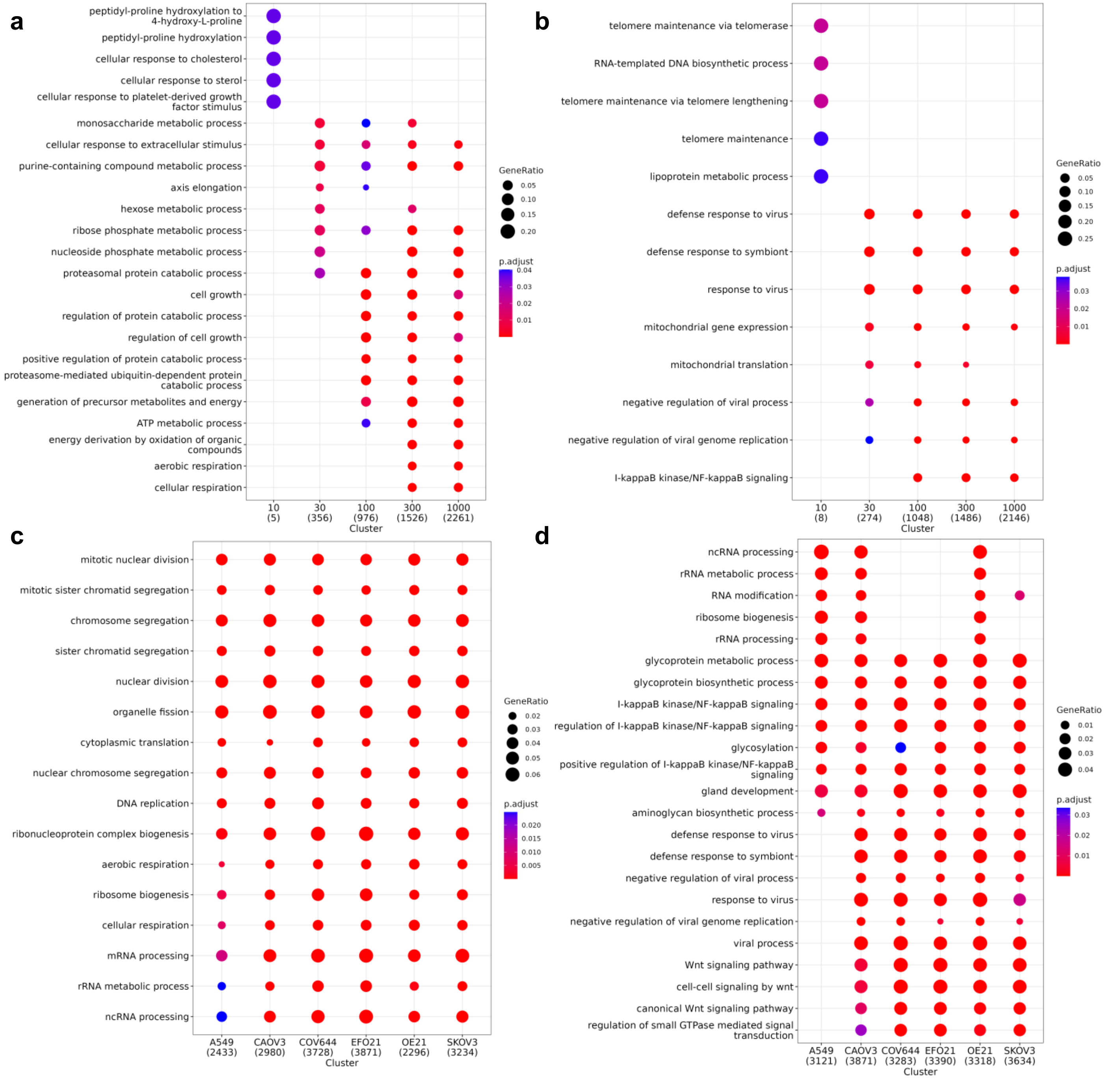


**Supplementary Figure 7.** Dot plots display enriched gene ontology biological process (GO:BP) terms for each group of genes, with dot size indicating the gene ratio (number of input genes associated with the term divided by the total number of input genes) and color indicating adjusted p-values. **(a-b)** Gene groups are genes with significantly decreased **(a)** or increased **(b)** RNA levels in Caov-3 cells treated with 10, 30, 100, 300 or 1000 nM STC-15 relative to DMSO-treated cells. **(c-d)** Gene groups are genes with significantly decreased **(c)** or increased **(d)** RNA levels in 6 cell lines treated with 1 μM STM3675 relative to DMSO-treated cells.

### Supplementary Figure 8.

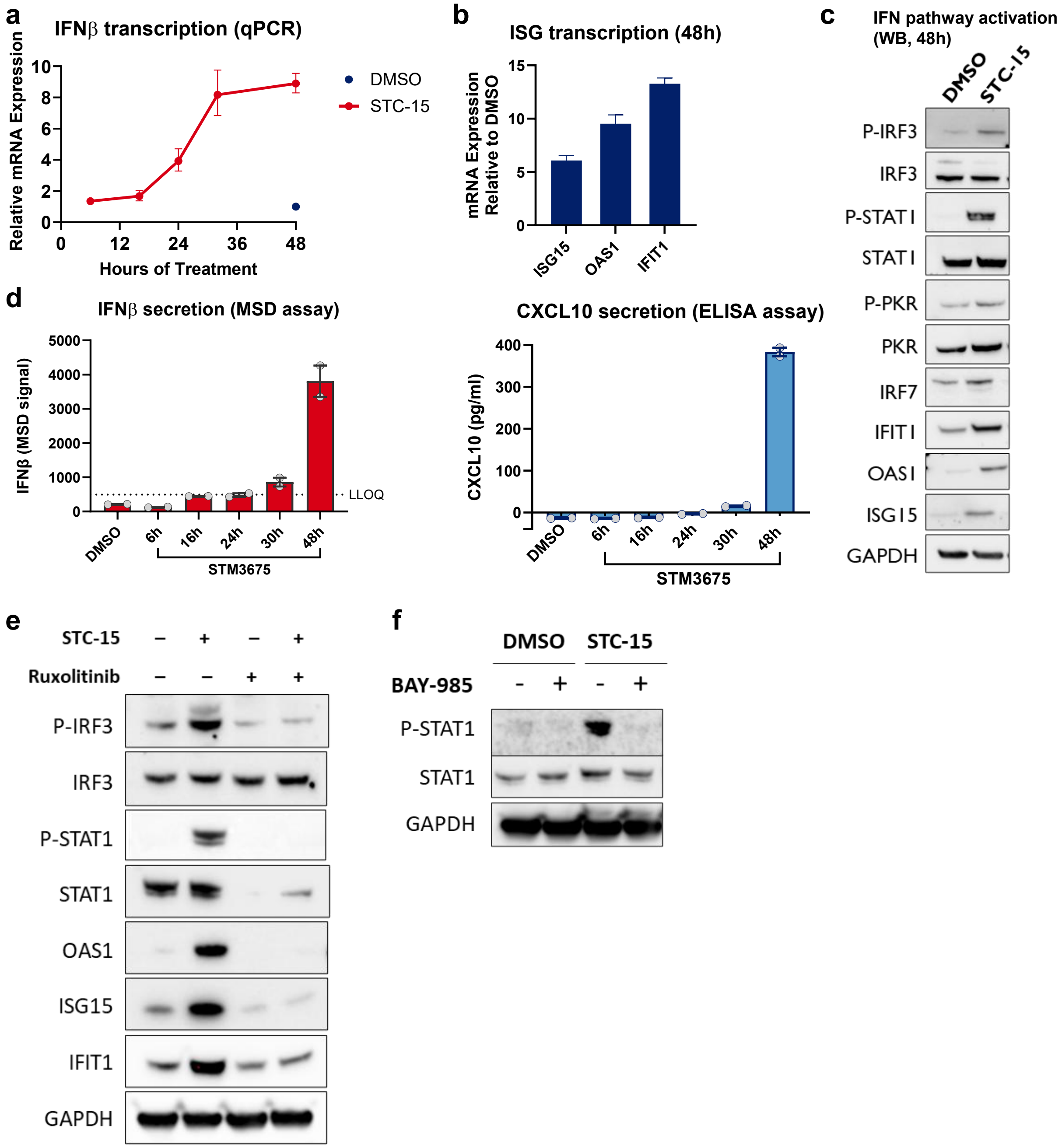


**Supplementary Figure 8.** Characterisation of IFN response in Caov3 cells. **(a-b)** Induction of IFNβ transcript over time **(a)** and selected ISG transcripts at 48 h **(b)** in Caov3 cells treated with 0.5 μM STC-15. Mean and range are presented for each timepoint. **(c)** A Western blot showing the activation of IFN signaling cascade in Caov3 cells treated with 0.5 μM STC-15 for 48 h. **(d)** Protein level of IFNβ (left) and CXCL10 (right) as measured in the growth medium of Caov3 cells treated with 2 μM STM3675 for the indicated times. Data are presented as mean ± SEM. **(e)** Western blot showing IFN signaling in Caov3 cells treated with 0.5 μM STC-15 for 48 h, with or without co-treatment with 1 μM Ruxolitinib, a JAK1/2 inhibitor. **(f)** Western blot showing IFN signaling in Caov3 cells treated with 1 μM STC-15 for 24 h, with or without co-treatment with 0.5 μM of BAY-985, a TBK1/IKKε inhibitor.

### Supplementary Figure 9.

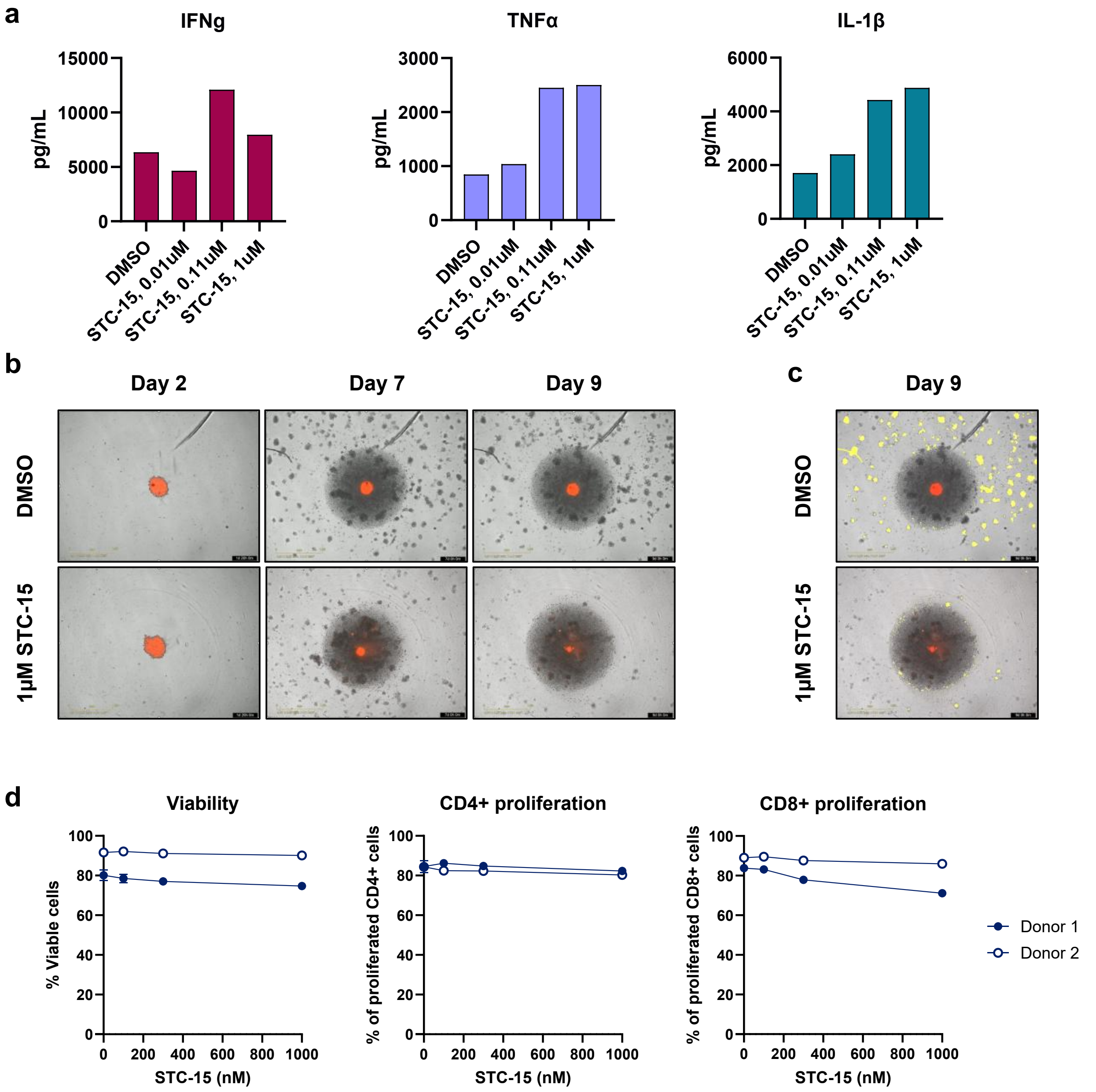


**Supplementary Figure 9.** Additional data from the co-culture system. **(a)** Secretion of pro-inflammatory cytokines in the co-culture experiment presented in Fig. 6A. Medium from 3 technical replicates was combined and assessed for the indicated cytokines using a Luminex assay. Data from one donor is presented and is representative of all three donors. **(b)** Representative IncuCyte brightfield images of SKOV3-NLR spheroids (red) and PBMC clusters (black) in different timepoints. **(c)** Quantification of PBMC migration into the centre of the well was achieved by applying a brightfield mask (yellow) over the PBMC clusters outside the main spheroid area and calculating the total mask area per image. **(d)** Plots showing flow cytometry assessment of cell viability and T-cell proliferation. PBMC from two donors were incubated with STC-15 at the indicated concentrations for 24 h and then stimulated with CD3/CD28 for additional 48 h prior to analysis. Data are presented as mean ± SD.

### Supplementary Figure 10.

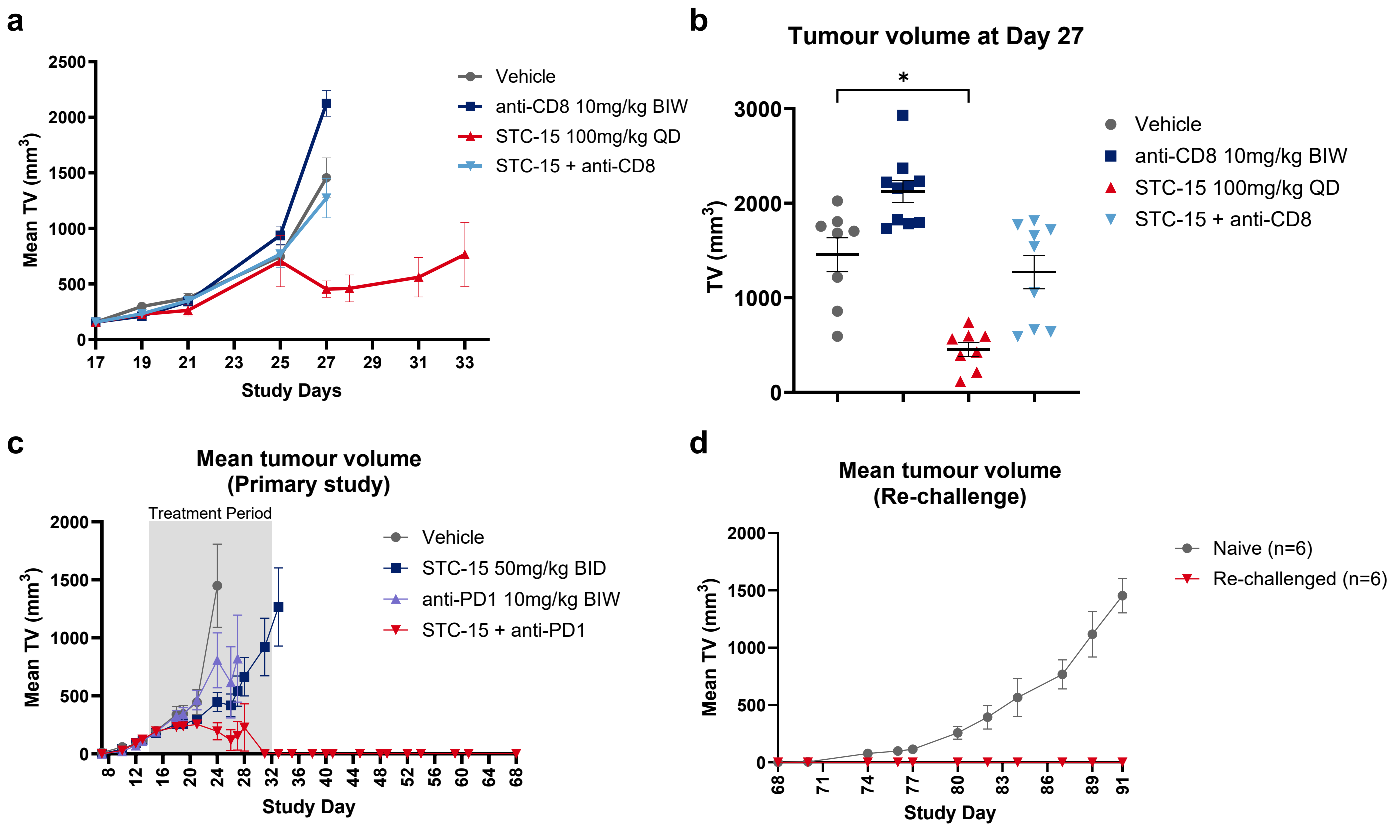


**Supplementary Figure 10.** STC-15 efficacy studies in the A20 syngeneic mouse model. **(a-b)** No STC-15 efficacy upon CD8+ cytotoxic T-cell depletion. Mean tumour volume over time **(a)** and individual tumour volume at Study Day 27 **(b)** of A20 sub-cutaneous tumours implanted immune competent host (BALB/c background). Mice (n= 10 / group) were treated with vehicle control or either STC-15, anti-CD8 depleting antibody or their combination at the indicated regimens. Data are presented as mean ± SEM. A group is shown while at least 50% of mice remain. **(c-d)** The combination of STC-15 and anti-PD1 leads to complete tumour regression (CR) and generates durable immune memory to reject newly implanted cells. Mean tumour volume over time **(c)** of A20 sub-cutaneous tumours implanted immune competent host (BALB/c background). Mice (n=7 / group) were treated with vehicle control or either STC-15, anti-PD-1 or their combination at the indicated regimens. Shaded area represents the treatment period (Day 14 to Day 32). A group is shown while at least 50% of mice remain. **(d)** Re-challenge: Following a treatment-free observation period of 27 days, 6 CR mice and a control group of 6 naïve mice were inoculated at Day 61 with fresh A20 cells on the left flank. No further treatment was given. The plot depicts mean tumour volume over time in the two groups. All data are presented as mean ± SEM.

### **Supplementary Figure 11.** STC-15 *in vivo* efficacy in syngeneic mouse models (MC38 syngeneic model)

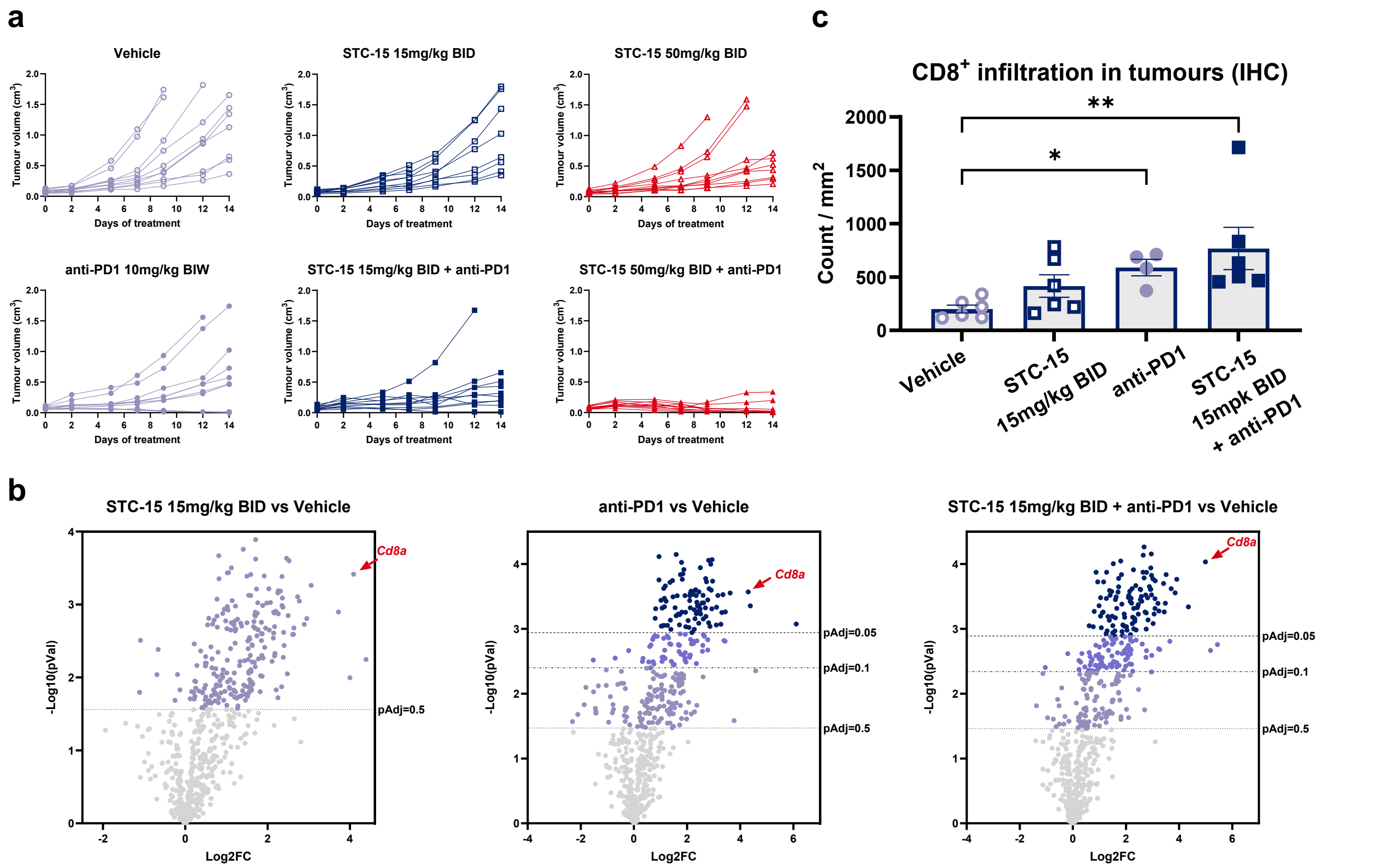


**Supplementary Figure 11.** Additional analyses of the MC38 study described in Fig. 7. **(a)** Volume of individual MC38 tumours. **(b)** Volcano plots showing differentially expressed genes in the Nanostring IO-360 dataset. Red arrows denoting the *CD8a* transcript in each comparison. **(c)** Number of CD8+ cytotoxic T-cells infiltrating the tumour tissue based on immunohistochemistry. Data are presented as mean ± SEM. Statistical analysis was performed using ANOVA testing with multiple comparisons. Each group was compared to Vehicle control. Statistically significant comparison are denoted by asterisks (*, $p<0.05$; **, $p<0.01$).
